# The role of the hypothalamus in cortical arousal and sleep homeostasis

**DOI:** 10.1101/2020.05.19.104521

**Authors:** Tomoko Yamagata, Martin C. Kahn, Merima Šabanović, Mathilde C.C. Guillaumin, Vincent van der Vinne, Yi-Ge Huang, Laura E. McKillop, Aarti Jagannath, Stuart N. Peirson, Edward O. Mann, Russell G. Foster, Vladyslav V. Vyazovskiy

## Abstract

Sleep and wakefulness are not simple homogenous all-or-none states, but instead are characterized by rich dynamics of brain activity across many temporal and spatial scales. Rapid global state transitions between waking and sleeping are believed to be controlled by hypothalamic circuits, but the contribution of the hypothalamus to within-state changes of sleep and wake “intensity” remains largely unexplored. Here we show that stimulation of inhibitory neurons in the preoptic hypothalamus does not merely trigger awakening from sleep, but the resulting awake state is also characterized by increased cortical activity. This activation is associated with a faster build-up of sleep pressure, proportional to the arousal level. These findings show that hypothalamic systems thought to exclusively control global state switching, also regulate within-state “intensity”, which we propose as a key intrinsic variable in shaping the architecture of sleep/wake states across the 24h day.

## Introduction

Interspecies variation in the daily amount of sleep is strongly influenced by genetics factors (Allada and Siegel, 2008). However, individuals also possess a striking ability to adapt the timing and duration of sleep in response to a variety of intrinsic and extrinsic factors (Ungurean et al., 2020). The key regulators of “adaptive sleep architecture” are: (i) homeostatic sleep need; (ii) the endogenous circadian clock; and (iii) the necessity to satisfy other physiological and behavioral needs such as feeding or the avoidance of danger (Eban-Rothschild et al., 2018). It is unknown how and in what form these numerous signals are integrated within the neural circuitry that generates the rapid and stable transitions between sleep and wake states.

Brain state switching has been the main focus of circuit-oriented sleep research for decades. Early studies identified the preoptic hypothalamus as a primary candidate for a hypothesized “key sleep center” (Economo, 1930; Nauta, 1946; Sallanon et al., 1989), and subsequent studies confirmed the existence of sleep-active neurons in the ventral lateral and medial preoptic area (VLPO and MPO) of the hypothalamus (McGinty and Szymusiak, 2001; Sherin et al., 1996; Szymusiak and McGinty, 2008). Combined with the findings that orexin/hypocretin neurons are necessary to maintain wakefulness (Chemelli et al., 1999; Lin et al., 1999), a model was proposed in which the sleep/wake-promoting circuitries function as a flip-flop switch (Saper et al., 2001). This model was able to account for rapid and complete transitions between sleep and wakefulness, and for preventing state instability (Mochizuki et al., 2004) or an occurrence of mixed, hybrid states of vigilance (Mahowald et al., 2011). However, over the last decade, our knowledge of subcortical brain nuclei that control sleep has expanded steadily, leading to the identification of functional specialization within the sleep-control network, and in parallel, highlighting a previously underappreciated complexity (Chung et al., 2017; Herrera et al., 2016; Kroeger et al., 2018; Liu and Dan, 2019; Liu et al., 2020; Ma et al., 2019; Oishi et al., 2017; Weber et al., 2018; Zhang et al., 2015; Zhong et al., 2019).

A key question to emerge is how signals regulating sleep architecture are represented and integrated in hypothalamic state-switching circuitries to ultimately maximize ecological fitness (Eban-Rothschild et al., 2017). Although sleep homeostasis has been considered an important factor influencing sleep/wake transitions (Alam et al., 2014; Carter et al., 2009; Cirelli et al., 1995; Saper et al., 2010), relatively few studies have addressed whether and how sleep/wake controlling brain areas overlap with those involved in homeostatic sleep regulation (Ma et al., 2019). Furthermore, while homeostatic sleep pressure, measured as electroencephalogram (EEG) slow-wave activity (SWA), builds up as a function of global wake duration, it is also locally regulated by specific activities during wakefulness (Krueger et al., 2013; Vyazovskiy and Harris, 2013). The property of sleep and wake as brain states with flexible intensities on a global and local level suggests an additional complexity which is difficult to reconcile with a simple flip/flop switch model. For example, there is evidence to suggest that wake “intensity” contributes to the build-up of global homeostatic sleep need (Fisher et al., 2016; Vassalli and Franken, 2017), and the balance between intrinsic and extrinsic arousal-promoting and sleep-promoting signals ultimately determines the probability of state switching (Eban-Rothschild et al., 2018; Lazarus et al., 2019).

Here, we investigate the role of the hypothalamus in the bidirectional interactions between sleep/wake switching and sleep homeostasis. We used the recently established paradigm of optogenetic activation of GAD2 neurons in the lateral preoptic area (LPO) of mice (Chung et al., 2017), Consistent with this study, we showed that optogenetic activation of the LPO led to rapid wake induction from sleep, but this effect was also observed when structures surrounding LPO were stimulated. Surprisingly, GAD2^LPO^ neuronal stimulation did not merely trigger wakefulness, but the awake state produced by this stimulation was characterized by increased cortical activation – the established measure of arousal (McGinley et al., 2015). In turn, subsequent sleep was associated with increased levels of EEG SWA, indicative of more intense sleep. Thus, our results demonstrate a dual role of hypothalamic circuitry, in regulating both the transitions between vigilance states and their intensity. We posit that such a “switch with a spring” mechanism can allow for both adaptive and reactive homeostasis of sleep and wakefulness.

## Results

### Stimulation of GAD2 neurons in the preoptic hypothalamus results in wakefulness

We employed a previously described protocol of optical stimulation of GAD2-expressing neurons in the LPO of the hypothalamus, which results in a rapid shift from sleep to wakefulness (Chung et al., 2017). To achieve the expression of ChR2, we injected AAV-DIO-ChR2-EYFP and implanted an optic fiber in the region of interest in GAD2-cre mice (n = 14) (Fig. 1A). Subsequent histology confirmed virus expression in a broad area including the preoptic area and adjacent areas. Specifically, in 8 out of the 14 mice that expressed the virus, the tip of the optic fiber was located in the LPO (LPO group), while in the remaining 6 mice the fiber tip was located within other neighboring hypothalamic areas posterior to the LPO (nonLPO group) (Fig. 1B)(Paxinos and Franklin, 2001). Since optical stimulation may affect the animal’s sleep and behavior due to local heating (Owen et al., 2019) or due to direct effects of light (Tyssowski and Gray, 2019), we also analyzed a cohort of GFP animals as a control group, which received the AAV-DIO-EGFP injection and were implanted with an optical fiber in the same area. EEG and EMG electrodes were also surgically implanted, as has been done previously (Guillaumin et al., 2018), to enable the identification of the sleep-wake state of the animal.

**Figure 1.**
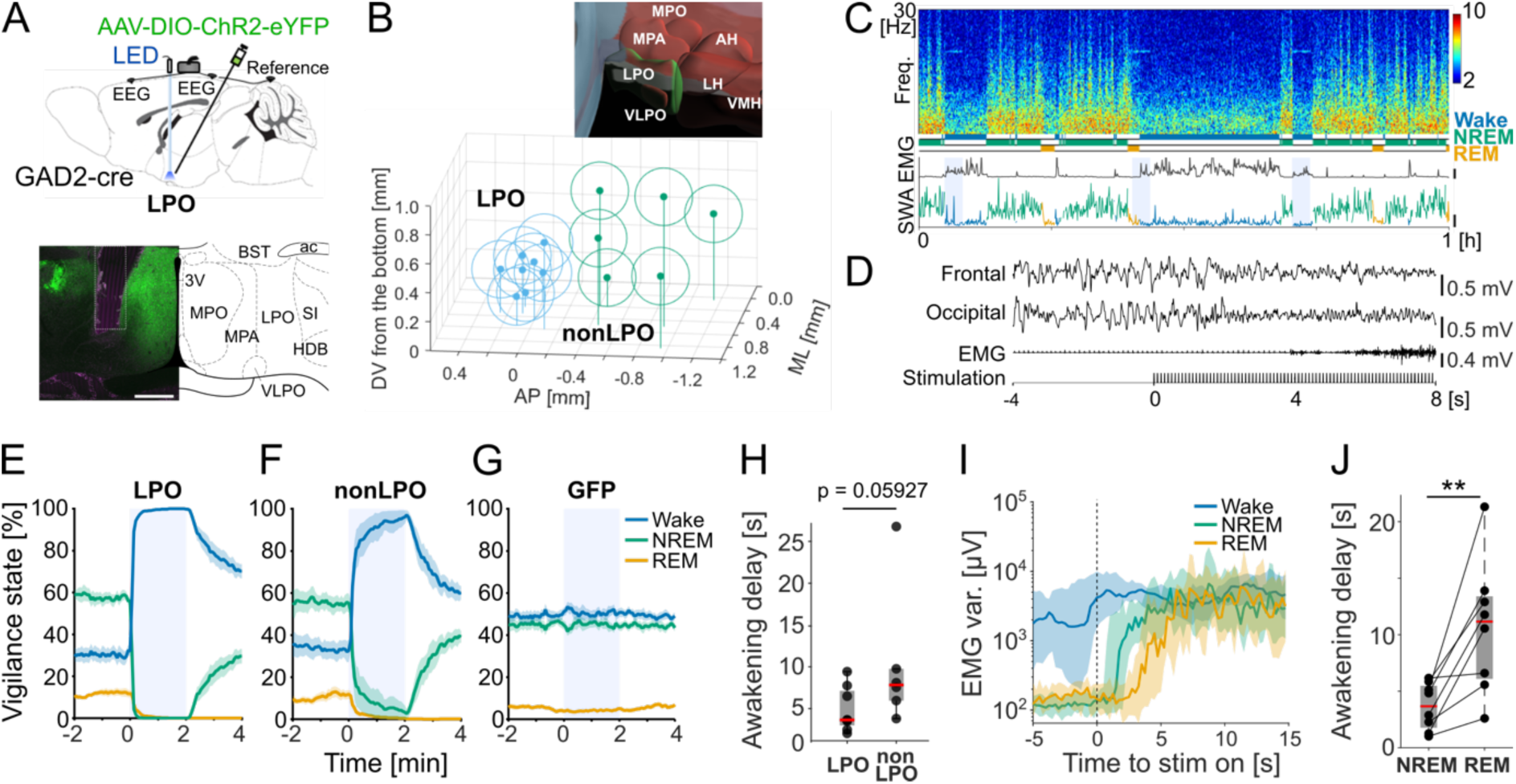
Optical activation of GAD2 neurons in the LPO and surrounding hypothalamic regions induces rapid awakenings. A) Top: Schematic diagram of the implant. Bottom: Representative brain section and corresponding brain areas from Paxinos and Franklin, the mouse brain atlas (2001). Scale bar, 500 *µ*m. B) Schematic of the optic fiber tip locations in all animals with ChR2 expression. The center of fiber tip is shown as dot, the stimulation coverage areas estimated based on a fiber diameter (400 *µ*m) are shown as circles for individual animals. Blue: LPO group, Green: nonLPO group. Top: Three-dimensional atlas of hypothalamic area, constructed by the Allen Brain Explorer (beta). C) Representative EEG spectrogram and the corresponding hypnogram, EMG and EEG SWA color-coded according to the state of vigilance. Scales, EMG: 5 mV; SWA: 1000 uV^2^/0.25Hz. Blue shade: photostimulation. Freq., frequency. D) EEG and EMG traces during a typical photostimulation trial in one individual animal. From top to bottom: frontal EEG, occipital EEG, EMG and the timing of photostimulation. E-G) Probability of wake, NREM and REM sleep before, during and after 2-min stimulation shown separately for the groups which received stimulation to the LPO area (n=8), nonLPO (n=6) and GFP controls (n=8). Blue shade, photostimulation. Mean values, SEM. H) Latency to awakening. Data points represent individual mice. Box represents 25/75 percentile and median values are in red. P-value is calculated by non-parametric two-sided Wilcoxon signed rank test. (LPO: n=8; nonLPO: n=6). I) Representative EMG variance profile (median ± 25/75 percentiles) averaged relative to the stimulation onset in one mouse. J) Latency to awakening for stimulations delivered during spontaneous NREM sleep and REM sleep in the LPO. Data points represent individual mice and box represents ± 25/75 percentiles across mice and red bar represents median. **p=0.007, two-sided Wilcoxon signed rank test (n=8). Abbreviations, 3V: third ventricle, ac: anterior commissure, AH: anterior hypothalamic area, BST: bed nucleus of stria terminalis, HDB: nucleus of the horizontal limb of the diagonal band, LH: lateral hypothalamic area, LPO: lateral preoptic area, MPA: medial preoptic area, MPO: medial preoptic nucleus; SI: substantia innominate, VLPO: ventrolateral preoptic nucleus, VMH: ventromedial hypothalamic nucleus.

In the first set of experiments, a 2-min period of light pulses at 10 Hz were delivered every 20 min throughout a full 24-h day, irrespective of the behavioral state (Fig. 1C, D and S1A). When stimulation was delivered during NREM sleep, the 2-min period of stimulation was completely dominated by wakefulness in GAD2-ChR2 animals implanted in both LPO (Fig. 1E, Suppl. Movie 1) and nonLPO locations (Fig. 1F), but not in the GFP controls (Fig. 1G). The average latency to awakening in the LPO group was on average 4.8 s (± 0.94 SEM, n=8) from the onset of stimulation (Fig. 1D, S1B) and was slightly longer in nonLPO group (10.3 ± 3.4 s, n=6, Fig. 1 H, p = 0.059, U=61, two-tailed Wilcoxon rank-sum test). Interestingly, when stimulation was delivered during REM sleep, awakenings were delayed compared to NREM sleep in LPO (12.58 ± 1.9 s; p = 0.007, Fig. 1 I, J) and in the nonLPO group (Fig. S1C). The latency to sleep upon cessation of stimulation was significantly longer if stimulation was performed when the animals were already awake (Fig S1D, Wake: 9.05 ± 0.59 min, NREM: 4.47 ± 0.42 min, REM: 5.22 ± 0.70 min, p<0.0001, one-way ANOVA). Thus, when applied during waking, stimulation of GAD2-neurons did not induce sleep but instead promoted wakefulness.

### Sleep/wake history does not influence the latency to awakening upon stimulation

We next hypothesized that elevated levels of sleep pressure decrease the probability of awakening upon photo-stimulation. To this end, mice were stimulated after 4-hour sleep deprivation (SD) or following an undisturbed period of sleep (Fig. 2A). As expected, SWA (EEG power density between 0.5-4 Hz) during NREM sleep in the 4-hour recovery period after SD was on average 65% higher than the SWA levels during the corresponding time period following undisturbed sleep (Fig. 2B). Thus, photostimulation during the sleep period after SD or after an undisturbed period of sleep are referred to as the “high-sleep pressure” (HSP), and “low-sleep pressure” (LSP) conditions, respectively. Unexpectedly, the animals still woke up almost immediately once stimulation was commenced (Fig. 2C) and the latency to awakening was indistinguishable between the HSP and LSP conditions (Fig. 2D, p>0.1, two-sided Wilcoxon signed rank test; Fig. S2A). This suggests that our stimulation protocol was too strong to be affected by physiological sleep pressure or that the targeted circuit is insensitive to homeostatic control of sleep and is therefore unaffected by the sleep/wake history. To further address these possibilities, we also compared the latency to sleep from stimulation offset and found that this was not significantly different between HSP and LSP conditions (Fig. S2B). The absence of an influence of sleep pressure was surprising as sleep intensity during NREM sleep following SD was initially more than twice as high as during sleep following no disturbances (Fig. S2C). Together, these results show that activation of a broad population of inhibitory neurons in the hypothalamus induces waking irrespective of homeostatic sleep pressure.

**Figure 2.**
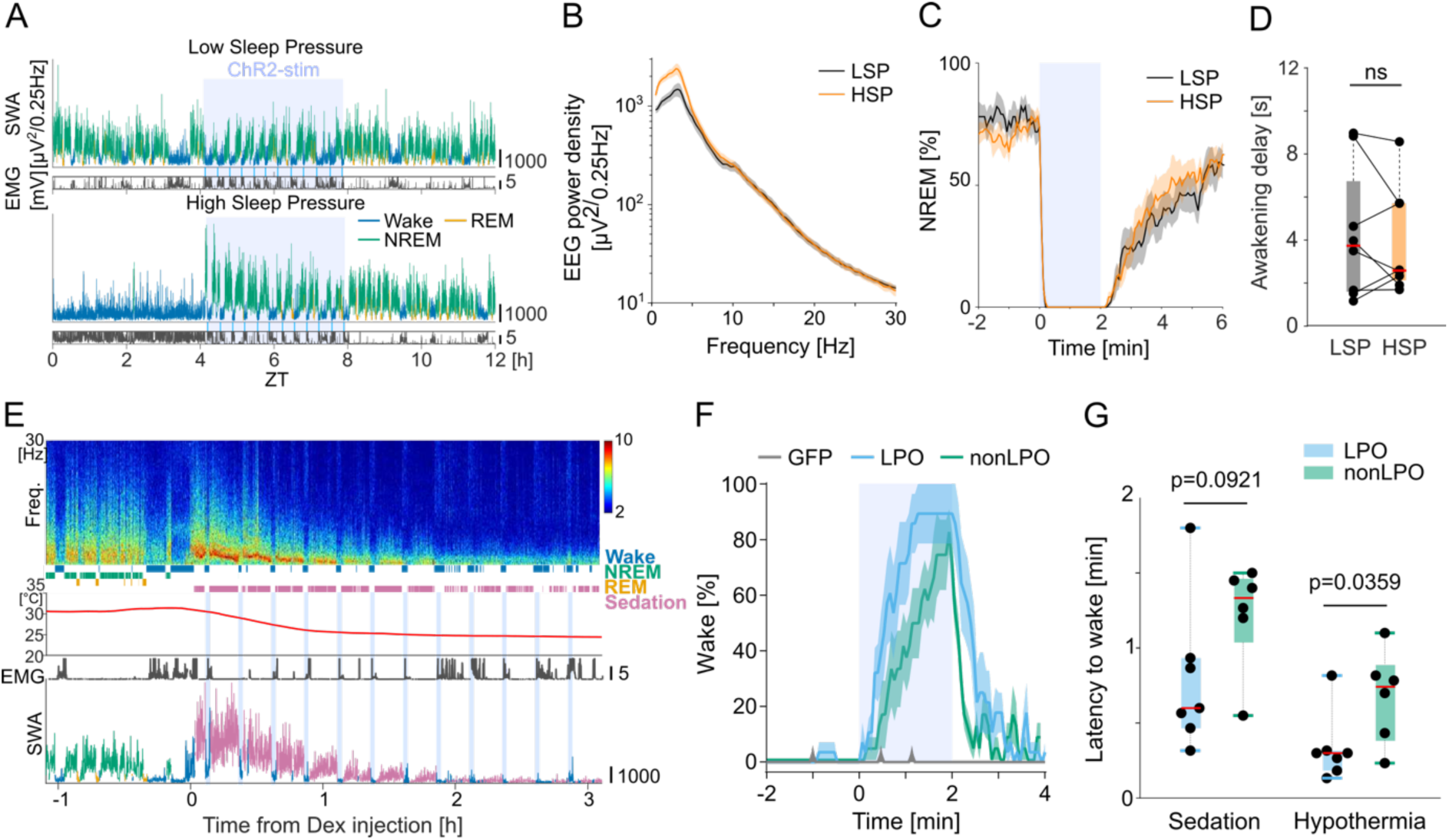
GAD2^LPO^ stimulation induces wakefulness independent of sleep pressure, and can reverse sedation. A) Representative 12-h profile of EEG SWA and EMG variance shown separately for “low” (top) and “high” (bottom) sleep pressure conditions. ZT: Zeitgeber time. B) Mean EEG power spectra during the first 4-h of recovery sleep after sleep deprivation (high sleep pressure = HSP) and the corresponding baseline interval when sleep pressure was low (LSP). Mean values, SEM. C) Percentage of NREM sleep occurrence before, during and after a 2-min photostimulation (shaded area) during high and low sleep pressure conditions. Mean values, SEM. D) Mean latency to awakening for stimulations delivered during high and low sleep pressure conditions. Data points represent individual mice and red bar represents median across mice. Box: ± 25/75 percentiles. A-D, LPO group, n=8. E) Representative frontal EEG spectrogram, hypnogram, peripheral body temperature, EMG and SWA color-coded according to the state of vigilance, from one mouse, before and after dexmedetomidine injection. Scales; EMG: 5 mV; SWA: 1000 *µ*V^2^/0.25Hz. F) Percentage of time spent awake before, during and after photostimulation (shaded area) during the first 1-hour interval after Dex injection, averaged for GFP (n=8), LPO (n=7) and nonLPO (n=6) animals. Mean values, SEM. Note GFP animals (grey) show rare occurrences of spontaneous wake. G) Latency to awakening from sedation after the onset of stimulation calculated separately for the initial sedation (average of 4 stimulation sessions during the 1st hour after Dex injection) and during late sedation under hypothermia (average of 4 stimulations delivered between 2-3 hours after Dex injection) in LPO (n=7) and nonLPO (n=6) animals. Red bar: median, box: ± 25/75 percentiles. P-value: unpaired t-test with Welch’s correction.

### Stimulation of GAD2 neurons awakens mice from Dexmedetomidine sedation

Although deep sleep after SD did not result in marked changes in the wake-promoting effects of hypothalamic GAD2 neurons stimulation, we hypothesized that sedation may produce a state where the effects of stimulation are ameliorated. Therefore, mice were sedated with dexmedetomidine (Dex) resulting in ‘deep sleep’-like EEG patterns and decrease of body temperature (Ma et al., 2019; Zhang et al., 2015), and delivered 10 Hz stimulation in 2 minute period as previously (Fig. 2E). Within minutes following the injection of Dex, the animals became immobile with an EEG dominated by high amplitude slow waves (Fig. S3A). As expected, peripheral body temperature and metabolic rate dropped rapidly and consistently (Fig. 2E, S3B, C). However, stimulation of GAD2 neurons during Dex sedation invariably resulted in awakening both in LPO and nonLPO groups (Fig. 2F). As expected, photostimulation had no effect in GFP control animals under sedation (Fig. 2F). Interestingly, upon stimulation onset, but before the animals woke up, a drop in EEG SWA relative to pre-stimulation levels was observed in the LPO group only (Fig. S3D, E). Furthermore, the latency to awakening from sedation tended to be shorter in LPO group as compared to nonLPO stimulated animals (Median 36 sec for LPO, 80 sec for nonLPO; p=0.092, unpaired t-test; Fig. 2F, G). Notably, although the latency to awakening was still substantially higher during stimulation under sedation as compared to spontaneous NREM sleep (Fig. S3F, NREM: 5.3 ± 1.6 s, SEM, Dex: 31.4 ± 5.5 s, n = 6, p<0.05, paired t-test), the animals showed largely normal behaviors after awakening, including exploration of novel objects (Suppl. Movie 2). After stimulation was terminated, the animals returned to a sedated state within a few minutes (Fig. 2F), and high amplitude SWA was re-established (Fig. 2E). Interestingly, the awakening invariably occurred even when stimulation was performed 2-3 hours after dexmedetomidine injection, when the peripheral body temperature dropped to nearly room temperature (Fig. 2G, S3G), and the latency to awakening was shorter as compared to trials performed during the first hour after injection (p<0.0021, two-way ANOVA, factor ‘0-1 hour vs 2-3 hour’ p=0.0021; post-hoc test, LPO: p=0.0273, nonLPO: p=0.0174).

### GAD2-photoactivation in LPO but not outside LPO generates excess sleep pressure

Wakefulness produced by photostimulation of GAD2 neurons typically outlasted stimulation both in the LPO and nonLPO regions by 4.69 ± 0.42 and 4.85 ± 0.54 min, respectively (p=0.82, Welch’s test). Given that the mice spent most of the time during stimulation sessions awake, and since a total of 72 stimulation sessions were delivered during the 24h period, we expected to observe a significant sleep loss. Indeed, the amount of sleep during the stimulation day was lower than the corresponding values during a baseline day in both LPO mice (86.3 ± 3 %) and nonLPO mice (86.4 ± 3.3 %), but not in the GFP group (100.4 ± 5.6 %, Fig. 3A, RM-ANOVA, p=0.035, t-tests against 100 % for LPO, p=0.002 (n=8), nonLPO, p=0.01 (n=6), GFP: p=0.7 (n=8)). However, while the suppression of NREM sleep during stimulation occurred in both LPO and nonLPO groups (Fig. 3B), and the overall amount was similarly reduced during the light period (Fig. 3A, S4A), we observed that the deficit in NREM sleep at the end of the 24h remained in the LPO stimulated group only (Fig. S4A). Furthermore, REM sleep was affected by stimulation to a greater extent than NREM sleep; the overall amount was reduced during the light period both in LPO and in nonLPO (Fig. 3A, S4A), but the proportion of REM sleep per total sleep time across 24 h was significantly reduced in the LPO group only (Fig. 3A, RM-ANOVA, p=0.012, post-hoc tests for LPO: p=0.048, nonLPO: p=0.098, GFP: p=0.067). These observed changes in NREM and REM sleep amount are in line with an interpretation where, because the repeated photoactivation was invariably followed by NREM sleep first, REM sleep was especially restricted by our experimental paradigm.

**Figure 3.**
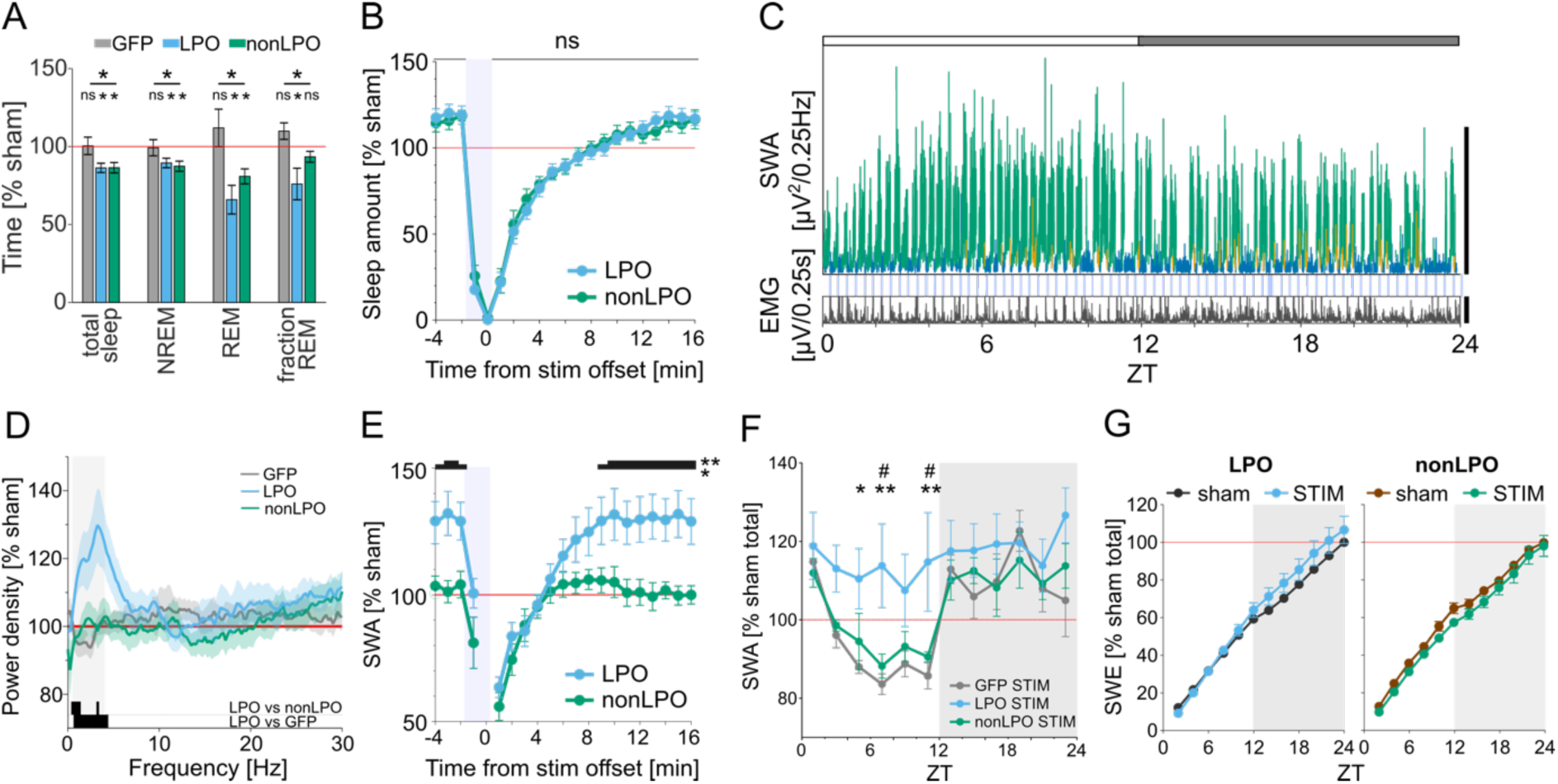
Photoactivation induces a rebound of SWA during NREM sleep in GAD2^LPO^ but not in GAD2^nonLPO^ animals. A) The effect of photostimulation on the total amount of vigilance states during the light period, shown as the % change relative to sham stimulation day. GFP: n=8; LPO: n=8; nonLPO: n=6. Stars above the lines on the top indicate significant difference in RM-ANOVA, and stars and ‘ns’ above plots indicate significance in t-tests against 100 %. *: p<0.05. Mean values, SEM. B) Amount of sleep during the 20-min window aligned to stimulation offset shown as percentage relative to the average sleep amount during sham stimulation day. All trials occurring during the light period are averaged for LPO (n=8) and nonLPO (n=6) animals. No significant difference was observed in sleep amount between the two groups in total (paired t-test, p=0.8167). Mean values, SEM. C) Distribution of EEG SWA and EMG during 24h in one individual animal that received stimulation to the LPO. SWA is color-coded according to vigilance state (green=NREM, blue = wake, orange = REM). Note the progressive increase in SWA across the light period. Scale bars for SWA, 5000 *µ*V^2^/0.25Hz; for EMG, 4mV. D) Relative spectral EEG power density in NREM sleep shown for the 10-Hz 24-hour stimulation condition during the light period, compared to sham condition. GFP (n = 7), LPO (n = 8), nonLPO (n = 6), Mean values, SEM. Black bars at bottom of figure denote frequency bins where the difference between experimental groups was significant (p<0.05). Shaded area denotes the SWA frequency range (0.5-4Hz). E) Time course of EEG over a 20-min window aligned to stimulation offset (time 0) during the light period. SWA is shown as percentage relative to SWA in sham stimulation day during the light period. *: p<0.05, **: p<0.01. Mean values, SEM. F) Time course of EEG SWA during NREM sleep across 24 hours on the day with photostimulations. SWA is plotted in 2-h intervals and represented as percentage of average NREM SWA during sham stimulation day (Mean values, SEM, GFP (n=7), LPO (n=8), nonLPO (n=6). *: p<0.05, **: p<0.01, Tukey’s multiple comparisons test after RM-ANOVA. *: LPO vs GFP, #: LPO vs nonLPO. G) Cumulative EEG slow-wave energy (SWE) across 24 hours. Mean values, SEM. LPO (n=8), nonLPO (n=6).

Sleep loss is compensated for not only by an increase in sleep time, but also by changes in sleep intensity, measured as the levels of EEG SWA. Strikingly, NREM sleep between stimulation sessions was characterized by an increase in EEG power selective to SWA over other frequencies in LPO group (Fig 3C, D, E, S4B, LPO: p = 0.0235, paired t-test), which was not observed in the nonLPO group or GFP controls (Fig. S4B). Since nonLPO and LPO mice lost a comparable amount of sleep during the light period, but the nonLPO group partially recovered the loss during the subsequent dark phase, we next assessed how NREM sleep SWA was distributed over the full 24h day. The time course of SWA across the day normally follows a U-shaped pattern with high SWA at light onset and decreased SWA in the middle of the 24h day (Fig. 3F). Unexpectedly, we found that this pattern was absent in the LPO group, which instead manifested, on average, a consistently elevated SWA, and in some individuals an increasing trend of SWA was present during the light phase (Fig. 3F, C). This altered daily distribution was not observed in the nonLPO or GFP control mice (Fig 3F, two-way ANOVA, p=0.0065; post-hoc Sidak’s multiple comparison test for LPO: p=0.0006, nonLPO: p>0.9999, GFP: 0.9804). Consistently, the increase in cumulative slow-wave energy (SWE) over the day in LPO mice was characterized by a steeper slope during the stimulation day compared to baseline or nonLPO mice (Fig. 3G, S4C, two-way ANOVA, p = 0.0406 for interaction of factors). Taken together, these results demonstrate that stimulation of the GAD2 neurons in LPO does not merely result in increased wakefulness, but that the induced wake state is characterized by a faster build-up of, or overall greater levels of sleep pressure needed to be compensated for.

### Stimulation of GAD2 neurons in the LPO results in increased levels of arousal

Previous studies suggest that the build-up of sleep pressure depends on the type of wake behaviors or levels of arousal, and not simply wake duration (Fisher et al., 2016; Suzuki et al., 2013; Vassalli and Franken, 2017). Therefore, we hypothesized that LPO and nonLPO stimulation differentially affects cortical activity within the awake state. To test this hypothesis, first EEG spectra were compared during wakefulness induced by stimulation with EEG spectra obtained after spontaneous awakenings during baseline, in both LPO and nonLPO mice. In both groups, stimulation-induced awakenings were associated with decreased frontal EEG power in slow wave range (Fig. 4A, B, general linear mixed model, p<0.001, F(97)=6.3 for main effect of frequency and p>0.5 for group and frequency x group, see Fig. 4 for post-hoc tests). However, stimulation-induced wakefulness was also characterized by a greater increase in occipital EEG theta-frequency power in the LPO group compared to the non-LPO group (Fig 4A-C, general linear mixed model, p<0.01, F(97)=1.91 for frequency x group, see Fig. 4B for post-hoc rank-sum tests in frontal and occipital EEG). Thus, stimulation of the LPO area did not simply trigger an awakening but the resulting wake state was also characterized by increased levels of arousal. We next hypothesized that altered wake “intensity” might account for the difference in sleep pressure between LPO and nonLPO mice (Fig. 3). In support of this hypothesis, across animals, a strong correlation was observed between the effects of stimulation on theta-frequency power during wake and the effect of stimulation on SWA during NREM sleep (Fig. 4C, Pearson’s R: 0.89). A relationship between the stimulation-induced decrease in SWA during wakefulness and SWA during NREM sleep was present but significantly weaker (Pearson’s R: 0.35, T(11)=3.72, p<0.01 for difference between correlation coefficients).

**Figure 4.**
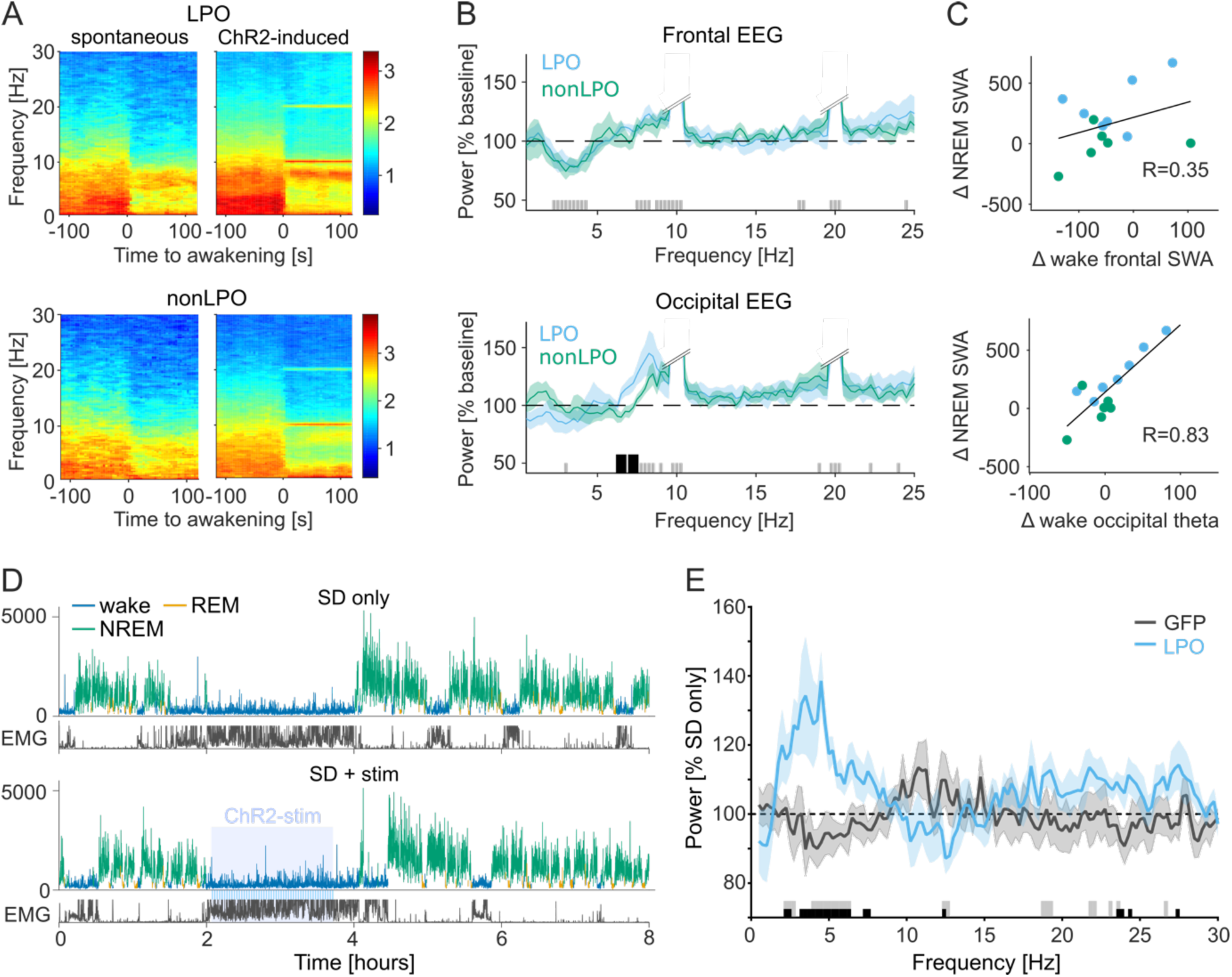
Effects of LPO and nonLPO stimulation on wakefulness and subsequent sleep. A) Average wake EEG spectrograms in occipital derivation in representative mice during spontaneous awakenings on a baseline day and stimulation-induced awakenings (Top: LPO, bottom: nonLPO). Time 0 corresponds to the onset of waking. B) The difference in average EEG spectral power density between spontaneous awakenings during baseline and photostimulation-induced awakenings. Bars at the bottom indicate significance (p<0.05) in non-parametric post-hoc tests; black: significant differences between conditions, grey: no differences between conditions but both conditions combined are different from 100%. Double slash: the cut-off of stimulation-induced artefacts. C) Correlation between stimulation-associated differences in wake theta power and NREM SWA. R=Pearson’s correlation coefficients. D) Example hypnogram illustrating the experimental design for stimulation during sleep deprivation (top: SD without stimulation (SD only), bottom: SD combined with 2-h photostimulation, shown as shaded area). SWA plotted for 4-s epochs is color-coded according to the state of vigilance. E) Effect of stimulation during wakefulness on EEG spectra during subsequent NREM sleep. The EEG power after SD + stim is calculated over the first 2-h of recovery sleep and represented as % of SD only condition in the LPO (n=6) and GFP (n=7) animals. Bars at the bottom denote frequency bins where the EEG power was significantly affected by stimulation (p<0.05 in multiple t-test, no correction for multiple comparisons); black: significant between GFP and LPO, grey: significant against 100% in LPO.

To directly address whether GAD2^LPO^ stimulation affects sleep homeostasis by modulating arousal during wakefulness, an additional experiment was performed. Photoactivation was performed during a 2h period while the animals were kept awake by providing novel objects (Fig. 4D, 20s trains at 10 Hz, delivered every 2 min). EEG SWA during subsequent NREM sleep was increased in LPO compared to the no stimulation condition (SD only), but not in GFP controls (Fig. 4E, LPO: p=0.0312, n=6, GFP: p=0.6875, n=7, Wilcoxon signed-rank test for mean SWA (0.5-4 Hz); LPO vs GFP: p=0.0290, two-tailed unpaired t-test for mean SWA). Thus, our data shows that GAD2^LPO^ activation during wakefulness increases levels of arousal and affects the build-up of sleep pressure.

Finally, to assess whether prolonged photoactivation could induce further increases in arousal, a subset of animals was exposed to an intense prolonged stimulation at 10 Hz for a 1-hour interval starting either at light or dark onset (Fig. S5). Unexpectedly, despite the animals being behaviorally awake, with brain activity and EMG indicated an unequivocal wake state, 3 out of 4 animals showed a marked decrease in peripheral body temperature (Fig. S5E), which in one case outlasted the stimulation period by approximately four hours (Fig. S5B).

### Cellular mechanism of GAD2^LPO^ mediated awakening

Recent studies on hypothalamic arousal circuitries raise two crucial questions regarding the cellular mechanisms underlying our observations. First, in some hypothalamic cells high frequency stimulation causes cells to enter a conduction block, which transforms optogenetic activation into *de facto* neuronal inhibition (Kroeger et al., 2018). Second, while the GAD2 promoter is thought to target mainly inhibitory cells, it has been shown that GAD2-cells can also be excitatory (Moffitt et al., 2018).

To assess whether conduction block only occurred at high stimulation frequencies, 1 Hz, 2 Hz and 5 Hz stimulation was sequentially applied to a subset of animals and compared to the awakening effect observed following stimulation frequencies of 10 Hz and 20 Hz (Fig. 5A-C). This showed that stimulation frequency had a pronounced effect on the probability to induce wakefulness (Fig. 5D, RM-ANOVA, p<0.01, n=7, see methods), and on the delay between stimulation onset and awakening (Fig. 5E, RM-ANOVA, p<0.001, n=7). However, frequencies below 10 Hz did not appear to result in a qualitatively different effect compared to stimulation at higher frequencies, instead, the wake promoting effect was stimulation-frequency dependent (Fig. 5A-C). In line with this, the latency to awakening was significantly influenced by the stimulation frequency (Fig. 5E, p<0.05 Wilcoxon signed-ranked test, n=7). Similarly, the probability that an awakening occurred within 2 minutes of stimulation onset was 75.0 % (± 5.1) at 2 Hz, increased to 90.6 % at 5 Hz, and peaked at 100% at 10 Hz (Fig. 5D, p<0.05 Wilcoxon signed-ranked test, n=7). Thus, while the possibility that some of the neurons transfected by ChR2 in our study entered conduction block cannot be excluded, the absence of a qualitatively different response following high-frequency stimulation contradicts this interpretation. Furthermore, the galaninergic neurons in the LPO, for which conduction block was previously demonstrated form only a small minority of LPO neurons (Kroeger et al., 2018).

**Figure 5.**
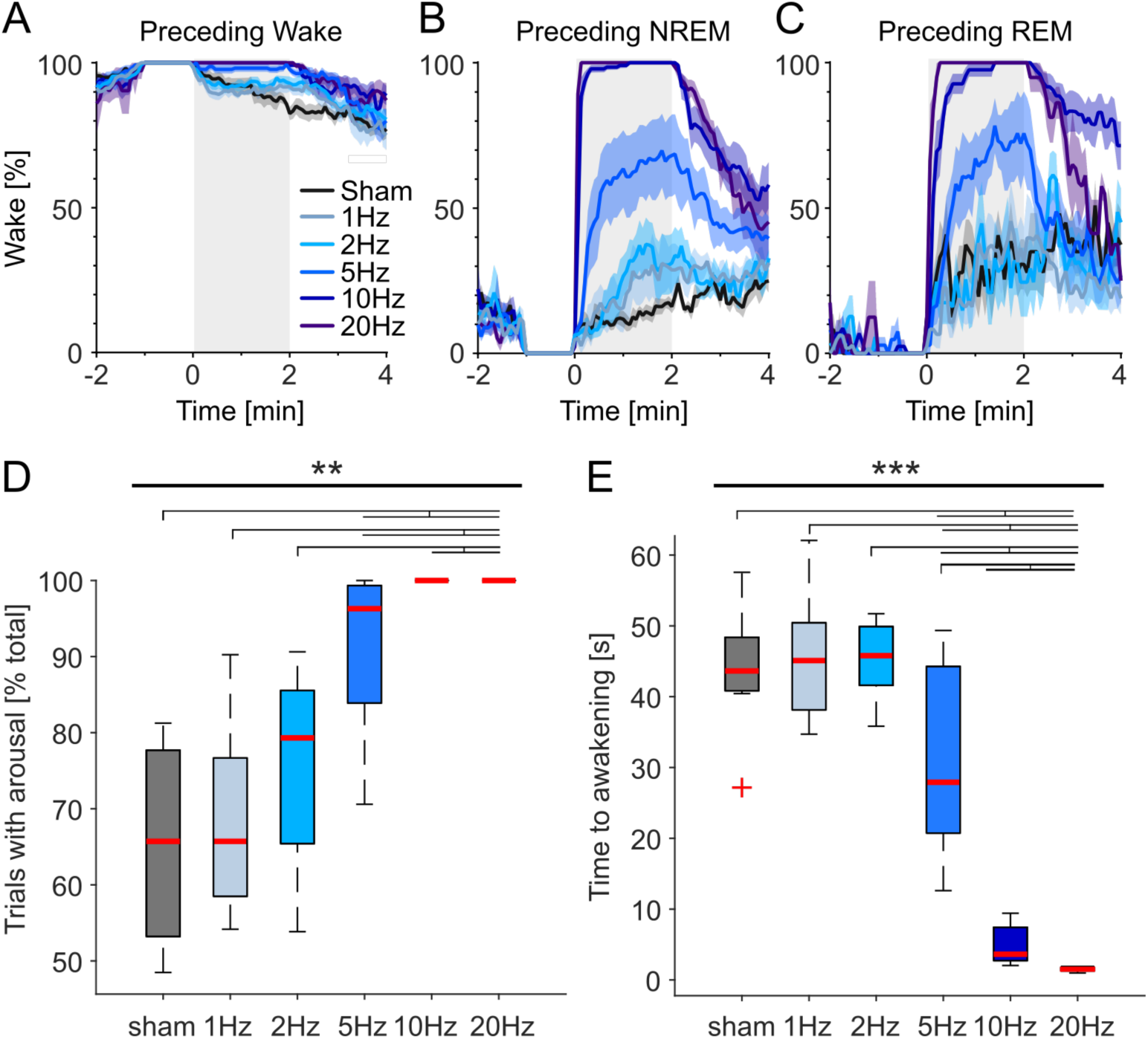
Low frequency stimulation has qualitatively similar effects on awakening latency as 10Hz stimulation. A-C) Effects of photostimulation (shaded area) on the amount of wakefulness. The trials are grouped according to the state of vigilance when stimulation started (wake and NREM sleep: last 1 minute, REM sleep: last 20 seconds before stimulation). Mean values, SEM; for sham, 1Hz, 2Hz, 5Hz, 10Hz: n=7 and for 20Hz: n=4. D) Proportion of NREM stimulation trials (± SEM), which result in an awakening within 2-min from stimulation onset. Stars (**) above bold line at the top denotes significance for RM-ANOVA (p<0.01). Lines above plots denote significant differences between conditions for pair-wise comparisons, Wilcoxon signed-rank test (p<0.05). E) Average latency to an awakening (± SEM) during those trials which resulted in an awakening. Stars (***) above bold line at the top denotes significance for RM-ANOVA (p<0.001). Lines above plots denote significant differences between conditions for pair-wise comparisons, Wilcoxon signed-rank test (p<0.05). Note that 20Hz stimulation only included 4 animals and was therefore not included in statistical comparisons.

The cellular substrate underlying the observed behavioral effects was further characterized by performing intracellular current clamp recordings from neurons in acute brain slices from GAD2^LPO^ mice. As this examination aimed to assess light responses in ChR2 expressing cells as well as their postsynaptic targets, cells in the LPO were targeted irrespective of ChR2 expression. As expected, a significant proportion of cells responded to negative current steps with hyperpolarization-induced sags and subsequent rebound low threshold spikes (LTS) and hyperpolarization-induced inwards currents (47 %, n = 85 cells, Fig. 6A, B, F). While almost all cells displayed spiking or subthreshold responses to light (90.6 %), the delay between stimulation and response distinguished cells that were directly activated by ChR2 (< 1 ms delay, 58%, Fig. 6B) from cells receiving synaptic inputs upon ChR2 activation. No differences were observed between LTS and non-LTS cells in the proportion of cells directly activated by ChR2 (Fig 6B, bottom panels; χ^2^(1,3) = 1.22, p = 0.75, Chi-square test). Next, we directly tested whether GAD2^LPO^ stimulation at 10 Hz is associated with conduction block and neuronal silencing. During 2 minutes of 10 Hz stimulation in putative ChR2 expressing cells, no signs of conduction block were found, and spike rates entrained to light pulses with high fidelity and independently of stimulation frequency (Fig. 6C, D; p = 0.70, one-way ANOVA). Indeed, average spike rates during the stimulation period were slightly higher than the stimulation frequency (Fig. 6E). These results are in line with our behavioral observation that effects of *in vivo* stimulation at 10 Hz are qualitatively similar to those at lower frequencies.

**Figure 6.**
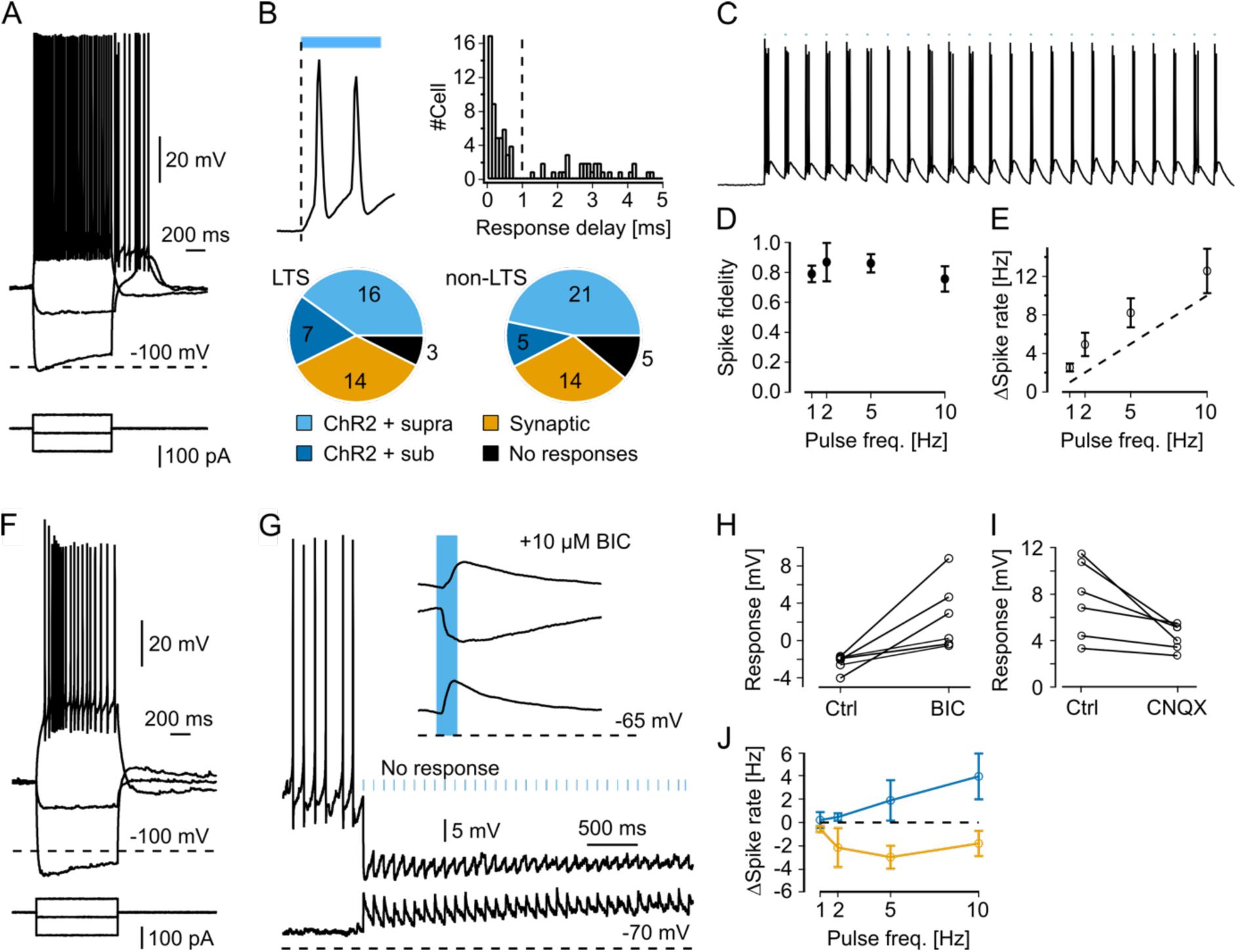
Single cell recordings in acute brain slices of GAD2^LPO^ mice. A) Representative electrophysiological characteristics of an LTS neuron in the preoptic area. B) Top left: Example of a membrane potential (Vm) response to ChR2 activation in a putative ChR2-positive cell (magnification of the trace shown on panel C). Note that depolarization starts immediately upon illumination. Top right: histogram of delays between optical stimulation and Vm response showing a bimodal distribution – responses with delay <1ms were classified as ChR2+, with delays >1 ms classified as synaptic responses. Bottom: Classification of cells according to ChR2 response properties. C) Example trace of a putative ChR2-positive cell responding to 10Hz stimulation. D) Spike fidelity of putative ChR2-positive neurons. Fidelity reflects the proportion of light pulses followed by a spike. E) Relationship between stimulation frequency and mean spike rate during stimulation in putative ChR2-positive neurons. F) Representative electrophysiological characteristics of a non-LTS neuron in the preoptic area. G) Representative traces and average evoked potentials (top right) of a ChR2-negative neuron, exhibiting depolarizing light responses at -70mV (bottom trace and bottom average evoked potential) and hyperpolarizing responses at a slightly depolarized Vm (upper trace, and middle in average evoked responses). The top average evoked potential shows the light-response at a depolarized membrane potential in the presence of bicuculline. Note the unmasking of excitatory responses. H) Change in average evoked responses (at depolarized Vm) in response to blocking of ionotropic GABA receptors with bicuculline. I) Change in average evoked responses (at resting Vm) in response to blockage of ionotropic glutamate receptors with CNQX. J) Changes in spontaneous spike rates (induced by injection of depolarizing currents) during 2 minutes of optical stimulation at 1 Hz, 2 Hz, 5 Hz, and 10 Hz. Yellow line represents putative ChR2-negative cells and blue line represents putative ChR2-positive cells.

Although the GAD2-Cre line was chosen to predominantly target ChR2 expression in inhibitory neurons, it has been suggested that GAD2 can also be expressed in excitatory neurons (Moffitt et al., 2018). To examine the postsynaptic effect of stimulating GAD2-ChR2 cells, the light responses to stimulation at 0.5 Hz were analyzed in putative non-ChR2 positive cells in the LPO (cells with subthreshold light responses and a delay to onset > 1 ms). To distinguish GABAergic from glutamatergic inputs, light-evoked responses were recorded at different membrane potentials (Fig. 6G). At rest, non-ChR2 neurons responded to light with small depolarizing potentials, which turned into hyperpolarizing potentials at higher membrane potentials (Fig. 6G). This suggests that light induces a strong GABAergic response with a weaker glutamatergic component. In line with this, pharmacological inhibition of GABA-A receptors (10 *µ*M Bicuculline) abolished the hyperpolarizing potentials and unmasked excitatory responses (Fig. 6H; n = 6, p = 0.02, paired *t*-test). Conversely, excitatory responses were sensitive to blocking ionotropic glutamate receptors with 10 *µ*M CNQX, although this effect was weaker (Fig. 6I; n = 6, p = 0.04, paired *t*-test). Strong inhibitory and weak excitatory currents suggested that GAD2^LPO^ stimulation had an overall inhibitory effect. This interpretation was tested by raising the membrane potential of ChR2-negative cells to promote spontaneous firing before delivering 2 minutes of light stimulation at 1, 2, 5 and 10 Hz, mimicking our in *vivo* protocol. The spontaneous firing rates in ChR2-negative cells did not significantly depend on stimulation frequency (p=0.43 RM-ANOVA) although spontaneous firing rates were suppressed by stimulation (T(35) = -3.58, p = 0.001, one-sample t-test against 0; Fig. 6J). Note that our sample size was however likely too small to adequately resolve frequency dependency. Conversely, putative ChR2-positive cells increased their firing rates in the same experimental paradigm (Fig. 6J, blue trace). Thus, the most parsimonious interpretation of our experiments *in vivo* and in brain slices is that the effects of GAD2^LPO^ stimulation reported in this study are driven largely by activating inhibitory neurons and are unaffected by conduction block.

## Discussion

The current study is the first to report that stimulation of inhibitory neurons in the lateral preoptic area of the hypothalamus does not simply trigger a transition from sleep to wakefulness but also increases the EEG indices of arousal. In turn, heightened arousal during the wake state leads to a compensatory increase in homeostatic sleep pressure, suggesting that the LPO not only controls execute global state switching but also increases within-state “intensity”. Moreover, these findings suggest wake intensity as a crucial parameter in the regulation of sleep. Based on these findings, we propose extending the prominent flip/flop switch model by adding a qualitative dimension to sleep-wake control, which can be best represented as an in-built “switch with a spring”. We posit that when wakefulness is especially intense, the “spring” becomes more compressed. As a result, once wakefulness ends, a stronger sleep rebound follows to release the “pressure”.

Our observation of rapid awakening upon the start of stimulation confirms earlier findings (Chung et al., 2017) and is consistent with the notion that only a subpopulation of preoptic neurons are sleep-active or sleep-promoting (Kroeger et al., 2018; Modirrousta et al., 2004). Previous studies sometimes tacitly assumed that the same hypothalamic areas are implicated in *both* sleep promotion *and* sleep homeostasis. An alternative possibility is that sleep propensity may be enhanced if some of the strong wake-promoting areas are inhibited, consistent with the idea that sleep represents a default state of a neural network or the whole organism (Bandarabadi et al., 2020; Krueger et al., 2013). Therefore, even if accumulation of sleep need, in some form, occurs across many distributed brain networks (Bridi et al., 2019; Bruning et al., 2019; Honda et al., 2018; Lazarus et al., 2019; Muheim et al., 2019; Noya et al., 2019; Shi and Ueda, 2018; Tatsuki et al., 2016; Williams and Naidoo, 2020), it is likely that state switching is initiated from a relatively limited set of brain circuits, which have the capacity to integrate sleep-wake history related signals with other ecological and homeostatic demands. Whilst the biological substrate of global sleep homeostasis remains unclear, the question of which brain areas are involved in encoding the time spent awake or asleep seems tractable.

Interestingly, our data suggest that while LPO and nonLPO stimulation resulted in comparable sleep loss, only LPO stimulation generates the enhanced sleep pressure manifesting as increased SWA during subsequent sleep. Furthermore, only LPO stimulation resulted in an increase in the EEG theta-frequency activity, an established marker of behavioral arousal (Kramis et al., 1975; Vassalli and Franken, 2017; Vyazovskiy and Tobler, 2005), during induced awakenings. Consistent with the observation that the build-up of sleep pressure is especially prominent during theta-rich wakefulness (Vassalli and Franken, 2017), we observed that the increase in EEG theta activity during LPO stimulation correlated strongly with the increase in SWA during subsequent sleep, while photoactivation of wake-promoting neurons during spontaneous wakefulness led to a further increase in sleep intensity.

The observed latency to awakening was longer when stimulation was delivered during REM sleep compared to NREM sleep. This observation argues against the possibility that our stimulation paradigm was either too strong or completely unphysiological. However, surprisingly, preceding sleep/wake history did not noticeably influence the latency to awakening induced by stimulation. Arguably, if a specific brain area is essential for sensing the levels of sleep need, it should show inertia in sleep/wake transition when the homeostatic sleep pressure is high. Consistent with this notion, sleep deprivation decreased the latency to awakening induced by 20 Hz stimulation delivered to orexin/hypocretin neurons (Carter et al., 2009). Therefore, our observation that sleep/wake history did not influence the probability of awakening induced by hypothalamic stimulation support the idea that neural substrates of sleep promotion and sleep homeostasis are, at least to some extent, independent.

An acknowledged limitation of our approach is that we likely targeted a heterogeneous population of neurons, as the hypothalamus harbors many different cell types (Chen et al., 2017). We would like to stress that it remains unclear how fine the spatial resolution should be in stimulation experiments such as ours, to obtain meaningful insights into cerebral substrates of global control of sleep and wakefulness. More relevant is that our chosen approach may have resulted in an activation of circuitries not directly, or not exclusively, related to sleep-wake control. For instance, LPO is adjacent to the lateral hypothalamus circuits which contain GABAergic neurons that promote a wide range of functions – from reward-seeking behaviors to feeding and behavioral arousal (Goldstein et al., 2018; Kosse et al., 2017; Nieh et al., 2016). Moreover, the preoptic area of the hypothalamus contains inhibitory neurons involved in parental behaviors and various homeostatic processes, such as thermoregulation or fluid homeostasis (McKinley et al., 2015; Wu et al., 2014; Zhao et al., 2017). Importantly, the LPO receives inputs from several limbic areas including the septum, subiculum, infralimbic cortex, and the brainstem (Chou et al., 2002), which can relay salient or aversive signals and therefore can function as an alarm system in response to important physiological drives or threats. In addition, a large population of GAD2 neurons in the LPO project to the parietal cortex and frontal cortex (Chung et al., 2017; Gritti et al., 1997) and it has been shown that selective photoactivation of prefrontally-projecting preoptic GAD2 neurons modestly, but significantly, promote wakefulness (Chung et al., 2017). Thus, the connectivity of the LPO and particularly its inhibitory neurons support the view that this area plays a central role in integrating essential physiological drives linked to the regulation of cortical arousal.

Our dexmedetomidine experiments showed that activation of GABAergic neurons in the hypothalamus under deep sedation is sufficient to produce arousal, consistent with the observations that sleep-active GAD neurons express α2A-adrenergic receptors (Modirrousta et al., 2004), and that selective ablation of galaninergic neurons in the LPO diminishes the effects of dexmedetomidine (Ma et al., 2019). Once again, we observed that LPO stimulation in sedated animals was more efficient in producing wakefulness and increasing cortical activation, as compared to nonLPO stimulation. Surprisingly, animals showed largely normal wake behavior when stimulated during sedation, even though their peripheral body temperature was reduced to values close to room temperature. Furthermore, our finding that continuous intense stimulation of the LPO can produce sustained wakefulness and hypothermia at the same time, further highlights that our understanding of the circuitry involved in vigilance states control and thermoregulation is incomplete (Kroeger et al., 2018; Ma et al., 2019; Zhao et al., 2017).

In conclusion, our study demonstrates that the preoptic area of the hypothalamus is not merely involved in global sleep/wake transitions but plays a more nuanced role in modulating within-state intensity and sleep homeostasis. Our data illustrates the important role of the hypothalamus in vigilance state control and provides an important step towards a more holistic characterization of neural substrates of sleep regulation.

## Experimental Procedures

### Animals

The experiments were performed in male and female Gad2-IRES-Cre mice (Jackson Laboratory 019022; B6N.Cg-*Gad2*^*tm2(cre)Zjh*^/J). Optogenetic manipulation experiments were performed on adult mice (2-9 months of age when surgery was performed, LPO 124 day ± 16 SEM, nonLPO: 107 ± 27, GFP 135 ± 14, one-way ANOVA, p=0.5879). Each group included at least 1 female mouse (LPO, n=1, nonLPO, n=2, GFP, n=2). Mice were housed under a 12-hour light-dark cycle (lights on 9:00 and off at 21:00) with food and water available *ad libitum*. All experimental procedures were performed under a UK Home Office Project License in accordance with Animal (Scientific Procedures) Act 1986 and the guideline of the University of Oxford.

### Viruses

AAV5-EF1a-DIO-ChR2/H134R-eYFP and AAV1-EF1a-DIO-eGFP were obtained from UNC vector core.

### Device and surgery

For chronic electroencephalogram (EEG) and electromyogram (EMG) recordings, we used custom-made headmounts composed of three stainless-steel screw electrodes (Fine Science Tools) and two stainless-steel wires attached to an 8-pin surface mount connector (8415-SM, Pinnacle Technology Inc., Kansas), as previously described (REF FISHER). All surgeries were performed using aseptic surgical technique and body temperature was monitored and maintained throughout surgical procedures. Analgesics were provided peri- and post-surgery (Buprenorphine and Meloxicam) and animals were monitored closely post-surgery. For *in vivo* optogenetic experiments, the animals were anesthetized with isoflurane (3-5% for induction and 1.5-2% for maintenance) and placed in a stereotaxic frame. Bregma and Lambda were adjusted to be horizontally aligned. The virus was loaded into a Hamilton syringe attached to a 32-gauge needle and injected slowly (50-100 nL/min) using an infuse/withdrawal pump aiming for the left lateral preoptic area (LPO: AP 0, ML 0.7 mm, DV 5.4 – 5.7 from Bregma) or left hypothalamic area posterior to LPO (nonLPO: AP -0.1, ML 0.7 mm, DV 5.4 – 5.7 from Bregma). All animals in all groups were injected with 400 nL of virus, except for 4 mice in the LPO group where injected with 150nL. For optical stimulation, either an optic fiber (400 um diameter, Doric Lenses Inc, Quebec) or a custom made optrode, consisting of an optic fiber glued with tungsten wires, was inserted at 0.2 mm above the virus injection site. For EEG recording, two screws were fixed into the skull above the right frontal cortex (AP +2, ML +2 from Bregma) and the right occipital cortex (AP -3.5, ML +2.5 from Bregma) and a reference screw was inserted into the skull above the cerebellum (−1.5 mm posterior from Lambda, ML 0). For EMG recording, two wire-electrodes were inserted into the left and right neck muscles. The optic fiber, screws and EMG electrodes were secured to the skull using non-transparent dental cement (Super-bond) to minimize the leaking of light during optical stimulation. The data from those animals in which either the implantation of the optical fiber was not successful (i.e., fiber penetrated through the hypothalamus) or the virus transfection was not successful were excluded from the analyses (2/10 in LPO group and 3/9 in nonLPO group).

### Electrophysiological recordings

4 to 7 weeks post virus injection, the animals were individually housed in a clear Plexiglas home cage, placed inside a sound-attenuated recording chamber (maximum 2 cages per chamber). After habituation to the home cage for at least 2 days, the EEG/EMG recording device mounted on the mouse skull was connected to flexible recording cables, and a patch cord was connected to the optic fiber implanted on the skull. Recordings started after at least 1 day of habituation to the cables. For data acquisition, we used a TDT RZ2 (Tucker-Davis Technologies) recording system. The EEG/EMG signals were digitized at 305 Hz and amplified with a PZ5 amplifier (Tucker-Davis Technologies) and filtered with a 128 Hz low-pass filter.

### Optogenetic manipulations

To examine the involvement of hypothalamic GAD2 neurons in vigilance state switching and homeostatic regulation, we applied blue LED stimulation (470 nm, 10.8 – 13.2mW at fiber tip) of 10 ms per pulse at various frequency and train durations. For stimulation across 24 hours, we stimulated at 20, 10, 5, 2 or 1 Hz, for a 2-minute train duration, with a 20-minute inter-trial interval with a jitter of 10% (i.e. 2-min stimulation every 18-22 minutes) (Fig. 1, 3, 4A-C, 5, S1, S4). In baseline day, no stimulation was given. Sham stimulation day (sham) is corresponding with a baseline day, but the timing of 2-min for sham stimulation was taken from 10 Hz stimulation. To examine the effects of sleep pressure on awakening latency (Fig. 2A-D), we compared effects of stimulation after 4-h of undisturbed sleep vs 4-hour of sleep deprivation (SD) starting from light onset (ZT0). In the high sleep pressure (HSP) condition, SD was performed by providing novel objects such as paper towels, wood pieces, Styrofoam and cardboard, and also by gentle cage tapping if necessary. At ZT4, all animals were left undisturbed and after 10-20 minutes (to allow the animals to fall asleep) photostimulation was started. Stimulation consisted, as above, in 10 Hz, 2-min long stimulation sessions delivered every 20 minutes across 4 hours (12 stimulation sessions in total). In the low sleep pressure (LSP) condition, the animals were undisturbed between ZT0-4 and stimulation was performed during the corresponding 4-hour period between ZT4-8, as in the high sleep pressure condition.

In the dexmedetomidine sedation experiment, dexmedetomidine (Dex), a selective alpha2-adrenoceptor agonist (DEXDOMITOR, Zoetis), was diluted with saline and injected s.c. (100 *µ*g/kg) at ∼ZT1. EEG, EMG and peripheral body temperature were continuously recorded before and after injection. Typically, within 2-3 minutes after Dex injection, the EEG started to show high amplitude, synchronized slow waves, and within approximately 5 minutes the animals became immobile. Once fully sedated (typically after 10 min) optical stimulation was commenced as above (10 Hz, 2-min long stimulation sessions) every 15 minute across 4 hours. Note that stimulation during sedation was more frequent than the previously used 20 minutes inter-trial intervals to examine more trials under sedation. For examining behavior during stimulation upon sedation (Suppl. Movie 2), 5-min stimulation sessions were performed every 15 minutes over a 2-hour period. All animals were injected with antisedan (ANTISEDAN, Zoetis) 4 hours after the Dex injection, which allowed recovery from sedation.

To examine the effect of stimulation during waking on subsequent recovery sleep (Fig. 4D, E), we applied trains of stimulation at 10 Hz for 20 seconds every 2 minutes during a 2-hour sleep deprivation protocol. Sleep deprivation was achieved by providing novel objects as above. Stimulation was stopped 10 minutes before the end of sleep deprivation (55 stimulation sessions over 2-h period in total). In a subset of animals, continuous stimulation at 10 Hz was performed for 1 hour, starting at ZT0 or ZT12.

For continuous stimulation (Fig. S5) we performed 10 Hz photostimulation for 1 hour starting at ZT0 for the light period condition and at ZT12 for the dark period condition.

### Peripheral body temperature and metabolic rates

In a subset of animals and wherever body temperature measures are reported (Fig. 2, S3, S5), peripheral body temperature was measured using infrared thermal imaging cameras Optris Xi 70 and Xi 80 (Optris, Berlin, Germany) attached to the ceiling of the recording chamber. The change in peripheral body temperature was calculated by taking the maximum value over 4 seconds and averaging over 30 minutes using a moving average. In two non-implanted mice in a separate experiment, metabolic rates were measured using indirect calorimetry, Calo (PhenoSys, Berlin, Germany) with a sampling rate of approximately 4 seconds (Fig. S3C). The animals were singly housed in standard IVC cages and O_2_ consumption, CO_2_ production and water vapor were measured. On the experimental day, after baseline metabolic rates were recorded between ZT0-3, the animals were taken out of the IVC cage and provided Dexmedetomidine (100 *µ*g/kg) diluted in saline by s.c. injection as above. Immediately after the injection, the animal was quickly moved back to the cage and the metabolic rates were monitored for the subsequent 3 hours. Reference air was analyzed for 1-minute intervals every 15 minutes and measurements during these intervals were excluded from the analyses.

### Vigilance state scoring

Vigilance states were scored manually by visual inspection of electrophysiological signals using SleepSign software (Kissei Comtec). For scoring, recordings were resampled at 256 Hz with custom MATLAB (MathWorks) scripts and the data format was transformed to edf format, for compatibility with SleepSign. After scoring, EEG power spectra were computed every 4-sec window by fast Fourier transformation (Hanning window, 0.25 Hz resolution).

### Histology

At the end of the experiment, mice were deeply anaesthetized and transcardially perfused with 0.1M PBS followed by 4% PFA in PBS. For fixation, brains were placed in 4% PFA for at least 24 hours and then placed in PBS with 0.05% sodium-azide. For sectioning, brains were placed in 30% sucrose in PBS for at least 3 days for cryoprotection and were then sectioned into 50 *µ*m coronal slices using a microtome. Slices were placed on glass slides and mounted in VECTASHIELD HardSet with DAPI to stain for DNA (H-1500, Vector Laboratories). Fluorescence images were taken using a confocal microscope (Olympus Fluoview FV1000).

### Slice recording

*Ex vivo* experiments were conducted 5-10 weeks after bilateral injection of AAV5-EF1a-DIO-ChR2/H134R-eYFP into LPO. Acute slice preparation was performed as described previously (Bartram et al., 2017). Briefly, mice were deeply anesthetized using isoflurane and rapidly dislocated and decapitated into ice cold carbogenated NMDG solution containing (in mM): 92 NMDG, 2.5 KCl, 1.25 NaH2PO4, 30 NaHCO3, 20 HEPES, 25 glucose, 5 Na-ascorbate, 2 thiourea and 3 Na-pyruvate, 10 MgSO4 and 0.5 CaCl2, titrated to pH 7.2-7.4 with HCl. Coronal slices (300 *µ*m) were cut using a vibratome (Leica Vibratome VT1200S) in ice cold NMDG solution. Slices including the LPO were recovered in NMDG solution for 10 min at 30-34 °C, and then placed into an interface chamber in carbogenated aCSF containing (in mM) 126 NaCl, 3.5 KCl, 1.25 NaH2PO4, 1 MgSO4, 2 CaCl2, 26 NaHCO3, and 10 glucose at pH 7.2-7.4 when bubbled with carbogen gas (95% O_2_, 5% CO_2_), at RT for 1-6 h, before being transferred to the recording chamber (aCSF at 32-34°C).

For whole-cell patch-clamp recordings, recording pipettes (standard borosilicate glass capillaries, 1.2 mm OD, 0.68 mm ID, WPI) of 3-8 MΩ resistance were filled with intracellular solution containing: 110 mM potassium gluconate, 10 mM HEPES, 4 mM NaCl, 0.2 mM EGTA, 2 mM MgATP, 0.3 mM GTP-NaCl, 8 mM phosphocreatine-bisodium, 20 μg/ml glycogen (Life-Tech, R0551), 0.5 U/μl recombinant RNase inhibitor (Takara, 2313A) and 4 mg/ml biocytin and adjusted to pH 7.2 with KOH (290-300 mOsmol l^-1^). This intracellular solution is modified to allow for single cell RNA sequencing (Cadwell et al., 2016; Fuzik et al., 2016).

Current-clamp recordings were carried out using an Axon Multiclamp 700B amplifier (Molecular Devices) low-pass filtered at 3 kHz and digitized using an Axon Digidata 1440A or Instrutech ITC-18 at 10 kHz. Data acquisition and stimulation protocols were controlled using Axon pClamp or custom-written procedures in IgorPro (WaveMetrics). The input resistance was determined by a hyperpolarizing 20 pA step for 300 ms in each sweep. To activate ChR2, we used blue light delivered by an LED through a 40x water immersion objective (Olympus, LUMPLFLN).

In order to sample both ChR2-expressing and non-expressing cells, we targeted cells in the LPO area without monitoring eYFP expression. Once a stable intracellular recording was obtained, we performed a step protocol to characterize basic electrophysiological properties of the cell. Subsequently, all cells were subjected to 1 or 2 minutes of ChR2 stimulation at 1 Hz. If the cell exhibited suprathreshold responses, it was subsequently subjected to stimulation at 2 Hz, 5 Hz and 10 Hz. If a cell did not exhibit suprathreshold ChR2-responses, we sought to quantify the net effect of light stimulation on spontaneous firing. Therefore, we injected a mild depolarizing current via the patch pipette to elicit spontaneous firing. Subsequently, we subjected the cell to 1 Hz, 2 Hz, 5 Hz, and 10 Hz stimulation for 1 or 2 minutes. Following these 4 stimulation protocols, we repeated stimulation at 1Hz before and after a 10-minute bath-application of CNQX (10 *µ*M, Sigma) or bicuculline (10 *µ*M, Sigma).

## Author Contributions

T.Y. and M.C.K. performed *in vivo* behavioral and electrophysiological experiments. T.Y., M.C.K., M.C.C.G. and V.V.V. performed sleep scoring. T.Y., M.C.K., M.S. and V.V.V. conducted behavioral and electrophysiological data analyses. T.Y. and Y-G.H. performed body temperature recordings and T.Y. analyzed the data. T.Y., V.v.d.V. and L.E.M. performed metabolic recordings and analyses. T.Y., M.C.K. and E.O.M. performed *in vitro* experiments and M.C.K. and E.O.M. did *in vitro* data analyses. T.Y., A.J., S.N.P., E.O.M., R.G.F. and V.V.V. designed experiments. T.Y., M.C.K. and V.V.V. primarily wrote the manuscript with input from all authors.

## Acknowledgments

This work was supported by Wellcome Trust Senior Investigator Award (106174/Z/14/Z), Wellcome Trust Strategic Award (098461/Z/12/Z), John Fell OUP Research Fund Grant (131/032), FP7-PEOPLE-CIG (PCIG11-GA-2012-322050), Royal Society (RG120466). We thank Prof. Denis Burdakov and Dr Christin Kosse for advice on experiments. We thank members of V.V.V.’s lab and SCNi for help with experiments and for comments on the manuscript. The authors have no competing interests.

---

**Supplemental Figure 1:**
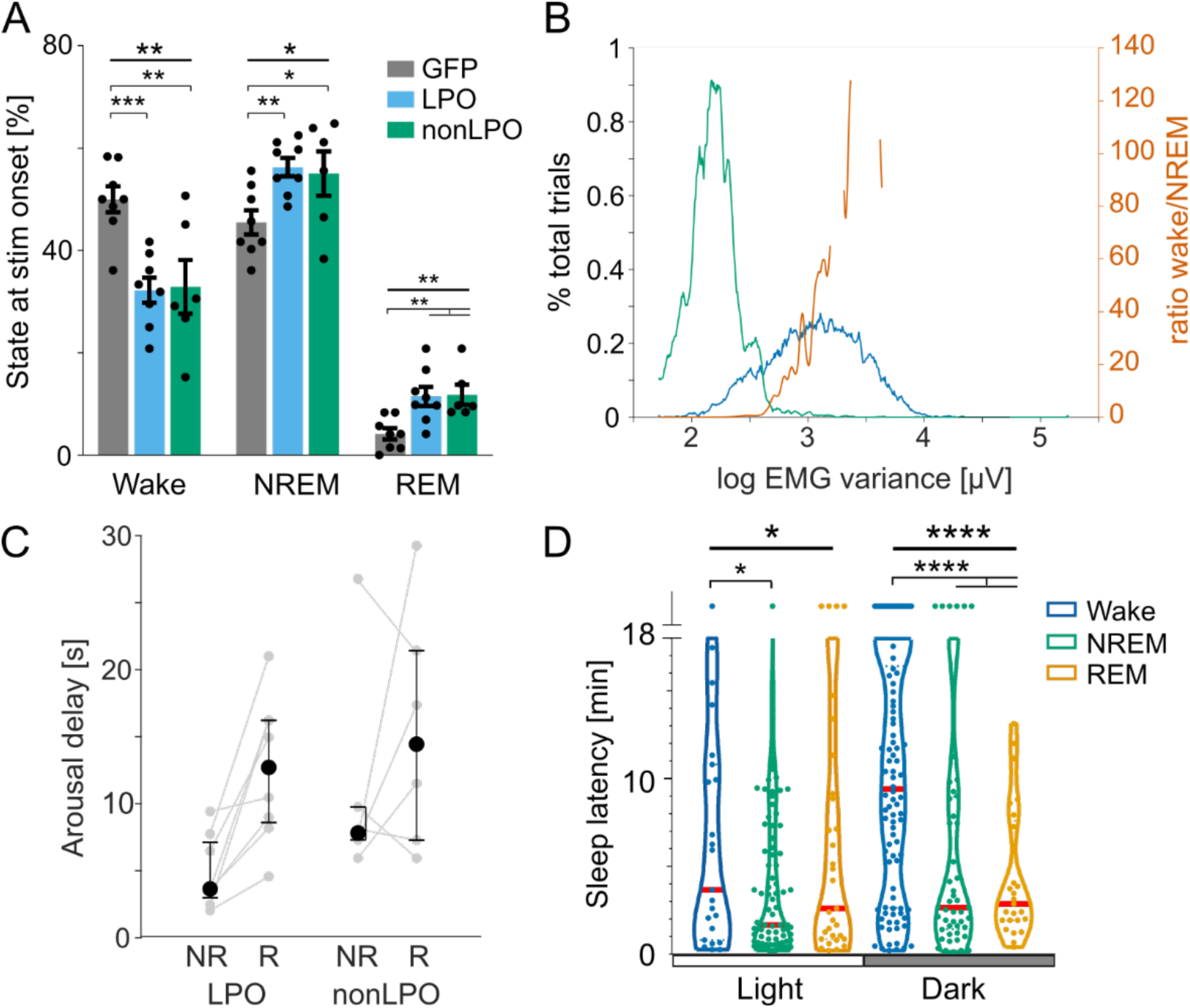
Effects of photostimulation. A) Percentage of photostimulation trials delivered during wake, NREM and REM sleep, averaged across animals for GFP (n = 8), LPO (n = 8) and nonLPO (n = 6) groups. *p<0.05, **p<0.01, ***p<0.001, RM-ANOVA and post-hoc uncorrected Fisher’s LSD. Error bar, SEM. B) Representative probability distribution of EMG variance across 24h, separated according to vigilance states (blue: wake, green: NREM), which were derived from visual scoring. For analysis of awakenings based on EMG variance, a threshold in EMG variance was derived from the ratio in probability between NREM and wake (orange line). C) Awakening delay after stimulation onset delivered during NREM and REM sleep in nonLPO and LPO animals. Grey points represent individual animals and black points show median ± 95% confidence interval across all mice. Mixed ANOVA, p=0.002 for factor ‘sleep state’; p=0.078 for ‘sleep state’ x ‘region’; p=0.079 for ‘region’. D) Latency to sleep after wake induced by photostimulation in LPO (n=8). Latency was calculated separately based on the vigilance state when stimulation was triggered. Top dots represent the number of cases in which animals did not go to sleep until the next photostimulation. *p<0.05, ***p<0.001, ****p<0.0001, one-way ANOVA and post-hoc uncorrected Fisher’s LSD. Red bar: median, dotted line violin plot: 25/75% across animals.

**Supplemental Figure 2:**
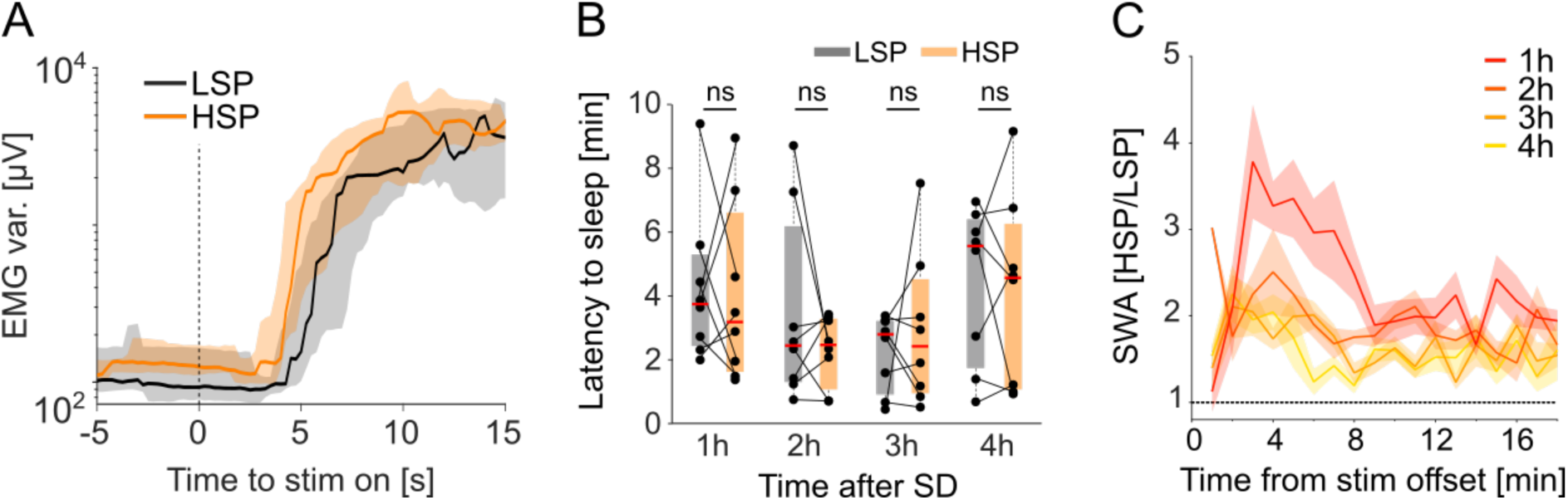
Photostimulation during high and low sleep pressure. A) Representative stimulus-aligned EMG variance (median ± 25/75 percentiles) in one mouse during high and low sleep pressure conditions. LSP: low sleep pressure condition, HSP: high sleep pressure condition. B) Latency to sleep after photostimulation during high and low sleep pressure conditions. Latencies were calculated in 1-h intervals during the 4-h recovery sleep after sleep deprivation. There were no significant differences between conditions at any time point (two-tailed paired t-tests). C) Time course of NREM sleep EEG SWA during the 18-minute interval between stimulations in the high sleep pressure condition. Values are plotted relative to mean SWA in the low sleep pressure condition. Dotted line at 1 means the same SWA in two conditions. Mean values, SEM.

**Supplemental Figure 3:**
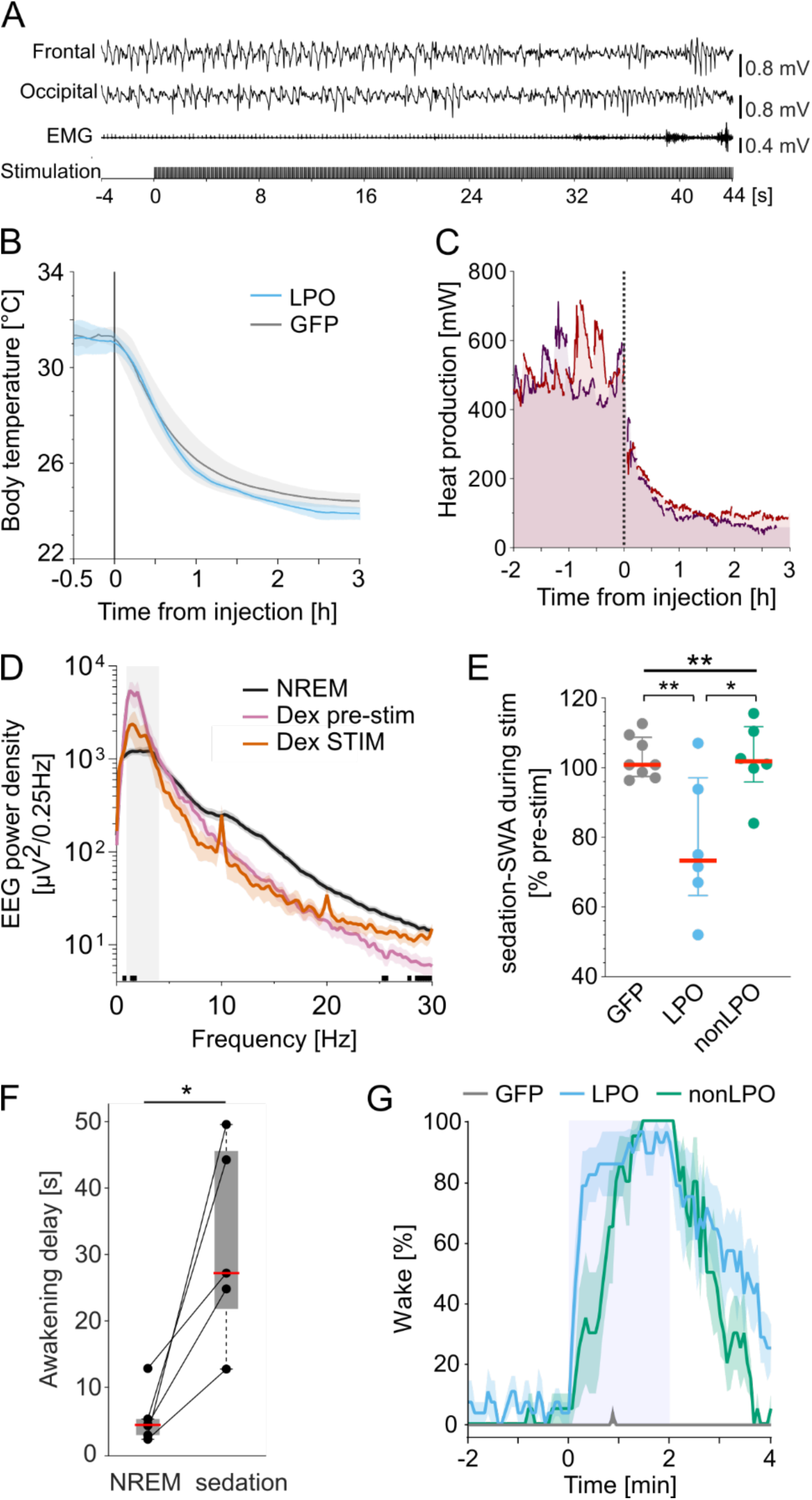
Photostimulation during dexmedetomidine sedation. A) Representative example of frontal and occipital EEGs and EMG traces before and during photo-stimulation in a representative animal injected with dexmedetomidine (Dex). B) Peripheral body temperature change during sedation with photostimulation in LPO and GFP control animals. Blue, LPO (n=4). Grey, GFP control (n=2). Stimulation was given in 2-min or 5-min long trains in LPO animals, and over 2-min long trains in GFP controls. Baseline temperature was obtained from average temperature during sleep before Dex injection between ZT 0-1. C) Metabolic rates calculated before and after Dex sedation. Intact Gad2-cre mice (n=2) were injected Dex at 0 (ZT 3) and metabolic rates were measured with indirect calorimetry. D) Average spectral power density of the frontal EEG during sedation in the LPO group (n=6). Pink: averaged over 2-min preceding photostimulations. Red: averaged over epochs after the onset of stimulation before awakening. Black, averaged 2-hour NREM sleep during ZT 1-3 in baseline day. Black bars at the bottom: p<0.05 in multiple t-test. Shaded area indicated SWA frequency band (0.5-4 Hz). E) EEG spectral power in the slow wave frequency range (SWA, 0.5-4 Hz) during 2-min 10 Hz stimulation in sedated animals, from GFP (n=8), LPO (n=6) and nonLPO (n=6) groups. Data are shown as percentage change from 2 minutes before stimulation. Wake states were excluded from sedation-SWA calculation. *: p<0.05, **: p<0.01 for one-way ANOVA and post-hoc Tukey’s multiple comparisons test. F) Latency to awakening for stimulations during NREM sleep or Dex-induced sedation. Datapoints represent individual mice and box represents median across mice (± 25/75 percentiles). *p<0.05, paired *t*-test (n=6). G) Probability of awakening from dexmedetomidine induced hypothermia. Average of 4 stimuli that were triggered between 2-3 hours after Dex injection. Blue, LPO (n=7). Green, nonLPO (n=6), Grey, GFP control (n=3).

**Supplemental Figure 4:**
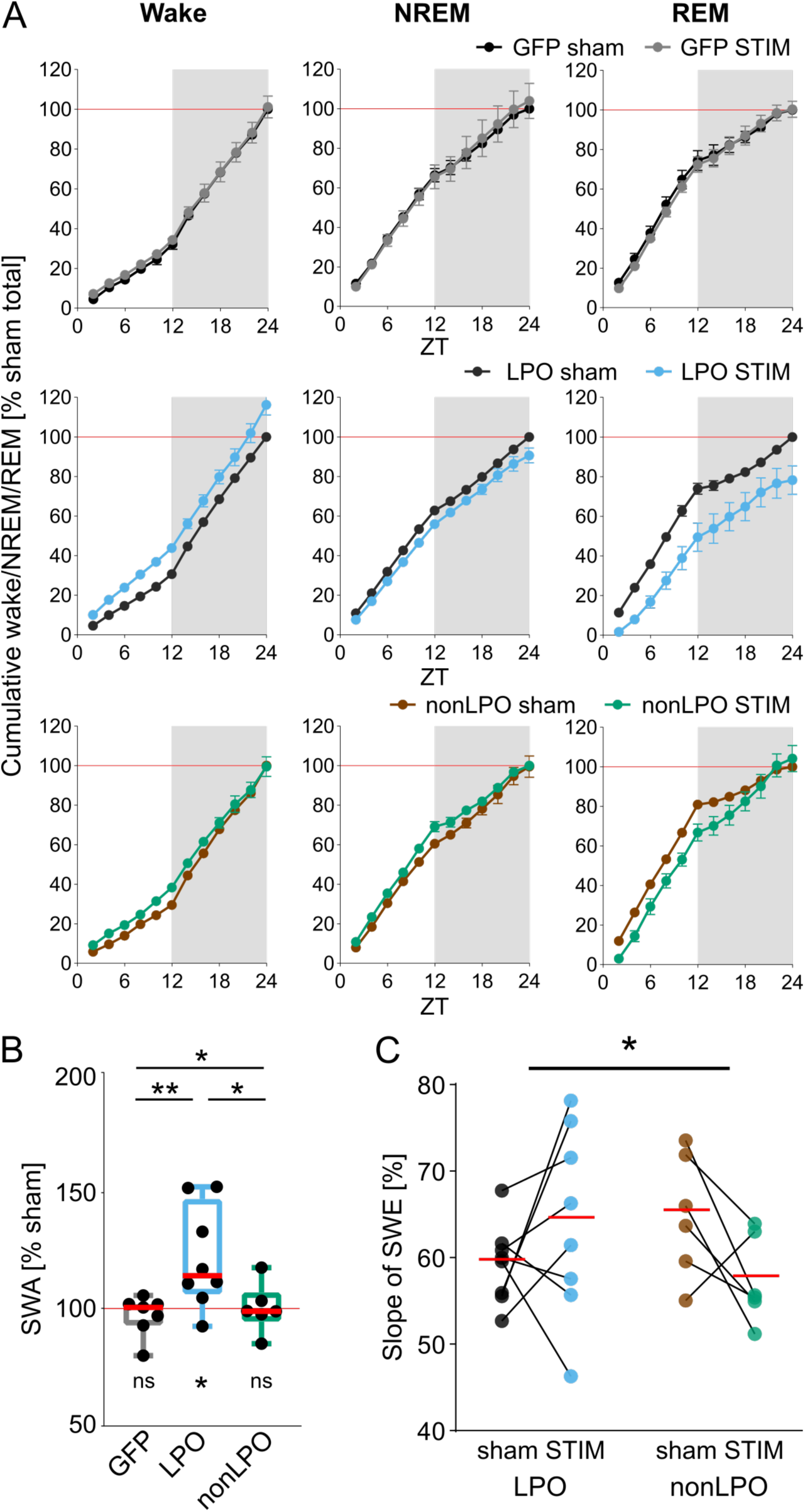
Effects of photostimulation on sleep and SWA. A) Cumulative amount of wake, NREM and REM sleep across a sham stimulation day and a day with photostimulation. The data are represented as percentage of the total amount of the corresponding state accumulated over the sham stimulation day (100%). Top panels: GFP-control (n = 8), middle panels: LPO (n = 8), bottom panels: nonLPO (n = 6). B) Frontal EEG-SWA during NREM sleep in a day with photostimulation shown as percentage of SWA in sham stimulation day. Comparison of relative EEG-SWA during NREM sleep in GFP, LPO and nonLPO groups. Top: *p<0.05, **p<0.01, one-way ANOVA and post-hoc uncorrected Fisher’s LSD test. Statistical results at the bottom were comparison of SWA between photostimulation day and sham stimulation day in each group for Wilcoxon signed-rank test; *: p<0.05, ns: no significance. C) Slope of EEG slow-wave energy (SWE) during the light period (Figure 3G). Red bar: median. *: p<0.05 for 2-way ANOVA for interaction between group and day.

**Supplemental Figure 5:**
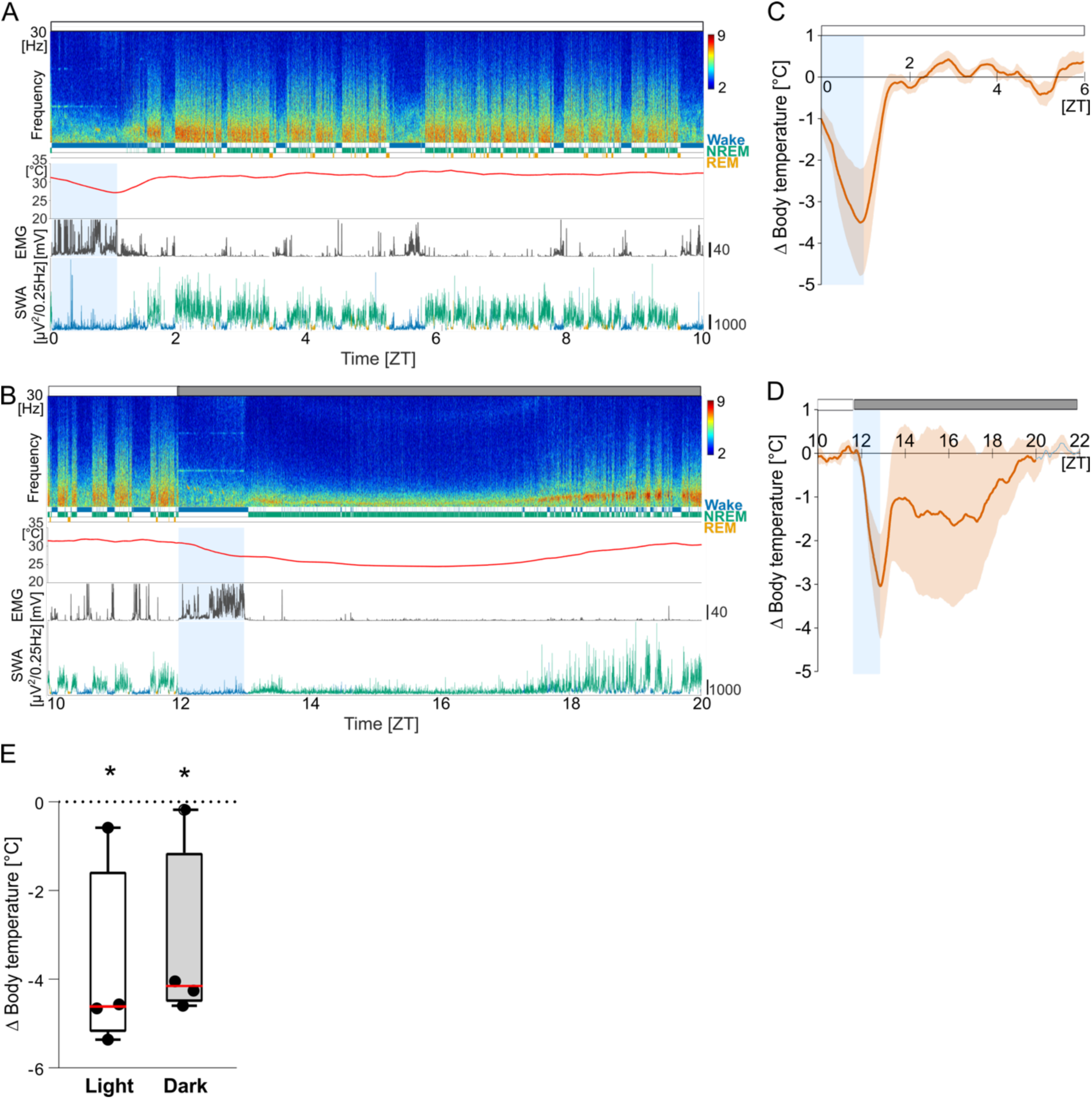
Photostimulation induced hypothermia. A, B) Representative spectrogram, hypnogram, peripheral body temperature, EMG and hypnogram-color-coded SWA during and after 1-hour 10 Hz stimulation starting at light onset (A) and dark onset (B). Data in A and B is from the same LPO animal recorded with an interval of 7-day. Note that, in A, the peripheral body temperature decreased consistently during stimulation but turned upward when stimulation was stopped and returned to pre-stimulation levels. In B, however, the temperature decrease persisted after stimulation was stopped. Scales; EMG: 40 mV; SWA: 1000 *µ*V^2^/0.25Hz. C, D) Average peripheral body temperature during and after 1-hour 10Hz stimulation compared to baseline levels in the LPO group (n=4). C) Stimulation at ZT0-1, D) at ZT12-13. Mean values, SEM. Shade, photostimulation. Baseline is calculated in each animal by averaging body temperature across 24 hours in baseline day. E) Change in minimum peripheral body temperature during photostimulation. Datapoints represent individual mice in the LPO group (n=4). Box represents 25/75 percentile. Red, median. *p<0.05, one-sample t-test.

**Supplemental Movie 1: Photoactivation of GAD2 neurons in LPO during NREM sleep induces wakefulness**.

This movie shows a representative response to a photostimulation trial. 30 s before and after photostimulation are shown, with photostimulation indicated by the black square in the bottom right corner. EMG activity is shown below to indicate movement. The movie runs at 2x speed.

**Supplemental Movie 2: Photoactivation of GAD2 neurons in LPO induces wakefulness in mice under Dexmedetomidine sedation**.

This movie shows the response of a Dexmedetomidine sedated mouse at different time points across the 5-minute photostimulation period. Specifically, video clips before stimulation (Pre stim), at the onset of stimulation (Stim on), 2-minutes into stimulation (During stim 2 min), 4 minutes into stimulation (During stim 4 min) and at the offset of stimulation (Stim off) are shown. The movie runs at 2x speed.

## References

Alam, M.A., Kumar, S., McGinty, D., Alam, M.N., and Szymusiak, R. (2014). Neuronal activity in the preoptic hypothalamus during sleep deprivation and recovery sleep. J Neurophysiol 111, 287–299.

Allada, R., and Siegel, J.M. (2008). Unearthing the phylogenetic roots of sleep. Curr Biol 18, R670–R679.

Bandarabadi, M., Vassalli, A., and Tafti, M. (2020). Sleep as a default state of cortical and subcortical networks. Curr Opin Physiol 15, 60–67.

Bartram, J., Kahn, M.C., Tuohy, S., Paulsen, O., Wilson, T., and Mann, E.O. (2017). Cortical Up states induce the selective weakening of subthreshold synaptic inputs. Nat Commun 8, 665.

Bridi, M.C.D., Zong, F.J., Min, X., Luo, N., Tran, T., Qiu, J., Severin, D., Zhang, X.T., Wang, G., Zhu, Z.J., et al. (2019). Daily Oscillation of the Excitation-Inhibition Balance in Visual Cortical Circuits. Neuron.

Bruning, F., Noya, S.B., Bange, T., Koutsouli, S., Rudolph, J.D., Tyagarajan, S.K., Cox, J., Mann, M., Brown, S.A., and Robles, M.S. (2019). Sleep-wake cycles drive daily dynamics of synaptic phosphorylation. Science 366.

Cadwell, C.R., Palasantza, A., Jiang, X., Berens, P., Deng, Q., Yilmaz, M., Reimer, J., Shen, S., Bethge, M., Tolias, K.F., et al. (2016). Electrophysiological, transcriptomic and morphologic profiling of single neurons using Patch-seq. Nat Biotechnol 34, 199–203.

Carter, M.E., Adamantidis, A., Ohtsu, H., Deisseroth, K., and de Lecea, L. (2009). Sleep homeostasis modulates hypocretin-mediated sleep-to-wake transitions. J Neurosci 29, 10939–10949.

Chemelli, R.M., Willie, J.T., Sinton, C.M., Elmquist, J.K., Scammell, T., Lee, C., Richardson, J.A., Williams, S.C., Xiong, Y., Kisanuki, Y., et al. (1999). Narcolepsy in orexin knockout mice: molecular genetics of sleep regulation. Cell 98, 437–451.

Chen, R., Wu, X., Jiang, L., and Zhang, Y. (2017). Single-Cell RNA-Seq Reveals Hypothalamic Cell Diversity. Cell Rep 18, 3227–3241.

Chou, T.C., Bjorkum, A.A., Gaus, S.E., Lu, J., Scammell, T.E., and Saper, C.B. (2002). Afferents to the ventrolateral preoptic nucleus. J Neurosci 22, 977–990.

Chung, S., Weber, F., Zhong, P., Tan, C.L., Nguyen, T.N., Beier, K.T., Hormann, N., Chang, W.C., Zhang, Z., Do, J.P., et al. (2017). Identification of preoptic sleep neurons using retrograde labelling and gene profiling. Nature 545, 477–481.

Cirelli, C., Pompeiano, M., and Tononi, G. (1995). Sleep deprivation and c-fos expression in the rat brain. J Sleep Res 4, 92–106.

Eban-Rothschild, A., Appelbaum, L., and de Lecea, L. (2018). Neuronal Mechanisms for Sleep/Wake Regulation and Modulatory Drive. Neuropsychopharmacology 43, 937–952.

Eban-Rothschild, A., Giardino, W.J., and de Lecea, L. (2017). To sleep or not to sleep: neuronal and ecological insights. Curr Opin Neurobiol 44, 132–138.

Economo, C.V. (1930). Sleep as a problem of localization. The Journal of Nervous and Mental Disease 71, 249–259.

Fisher, S.P., Cui, N., McKillop, L.E., Gemignani, J., Bannerman, D.M., Oliver, P.L., Peirson, S.N., and Vyazovskiy, V.V. (2016). Stereotypic wheel running decreases cortical activity in mice. Nat Commun 7, 13138.

Fuzik, J., Zeisel, A., Mate, Z., Calvigioni, D., Yanagawa, Y., Szabo, G., Linnarsson, S., and Harkany, T. (2016). Integration of electrophysiological recordings with single-cell RNA-seq data identifies neuronal subtypes. Nat Biotechnol 34, 175–183.

Goldstein, N., Levine, B.J., Loy, K.A., Duke, W.L., Meyerson, O.S., Jamnik, A.A., and Carter, M.E. (2018). Hypothalamic Neurons that Regulate Feeding Can Influence Sleep/Wake States Based on Homeostatic Need. Curr Biol 28, 3736–3747 e3733.

Gritti, I., Mainville, L., Mancia, M., and Jones, B.E. (1997). GABAergic and other noncholinergic basal forebrain neurons, together with cholinergic neurons, project to the mesocortex and isocortex in the rat. J Comp Neurol 383, 163–177.

Guillaumin, M.C.C., McKillop, L.E., Cui, N., Fisher, S.P., Foster, R.G., de Vos, M., Peirson, S.N., Achermann, P., and Vyazovskiy, V.V. (2018). Cortical region-specific sleep homeostasis in mice: effects of time of day and waking experience. Sleep 41.

Herrera, C.G., Cadavieco, M.C., Jego, S., Ponomarenko, A., Korotkova, T., and Adamantidis, A. (2016). Hypothalamic feedforward inhibition of thalamocortical network controls arousal and consciousness. Nat Neurosci 19, 290–298.

Honda, T., Fujiyama, T., Miyoshi, C., Ikkyu, A., Hotta-Hirashima, N., Kanno, S., Mizuno, S., Sugiyama, F., Takahashi, S., Funato, H., et al. (2018). A single phosphorylation site of SIK3 regulates daily sleep amounts and sleep need in mice. Proc Natl Acad Sci U S A 115, 10458–10463.

Kosse, C., Schöne, C., Bracey, E., and Burdakov, D. (2017). Orexin-driven GAD65 network of the lateral hypothalamus sets physical activity in mice. Proc Natl Acad Sci U S A 114, 4525–4530.

Kramis, R., Vanderwolf, C.H., and Bland, B.H. (1975). Two types of hippocampal rhythmical slow activity in both the rabbit and the rat: relations to behavior and effects of atropine, diethyl ether, urethane, and pentobarbital. Exp Neurol 49, 58–85.

Kroeger, D., Absi, G., Gagliardi, C., Bandaru, S.S., Madara, J.C., Ferrari, L.L., Arrigoni, E., Munzberg, H., Scammell, T.E., Saper, C.B., et al. (2018). Galanin neurons in the ventrolateral preoptic area promote sleep and heat loss in mice. Nat Commun 9, 4129.

Krueger, J.M., Huang, Y.H., Rector, D.M., and Buysse, D.J. (2013). Sleep: a synchrony of cell activity-driven small network states. Eur J Neurosci 38, 2199–2209.

Lazarus, M., Oishi, Y., Bjorness, T.E., and Greene, R.W. (2019). Gating and the Need for Sleep: Dissociable Effects of Adenosine A1 and A2A Receptors. Front Neurosci 13, 740.

Lin, L., Faraco, J., Li, R., Kadotani, H., Rogers, W., Lin, X., Qiu, X., de Jong, P.J., Nishino, S., and Mignot, E. (1999). The sleep disorder canine narcolepsy is caused by a mutation in the hypocretin (orexin) receptor 2 gene. Cell 98, 365–376.

Liu, D., and Dan, Y. (2019). A Motor Theory of Sleep-Wake Control: Arousal-Action Circuit. Annu Rev Neurosci 42, 27–46.

Liu, D., Li, W., Ma, C., Zheng, W., Yao, Y., Tso, C.F., Zhong, P., Chen, X., Song, J.H., Choi, W., et al. (2020). A common hub for sleep and motor control in the substantia nigra. Science 367, 440–445.

Ma, Y., Miracca, G., Yu, X., Harding, E.C., Miao, A., Yustos, R., Vyssotski, A.L., Franks, N.P., and Wisden, W. (2019). Galanin Neurons Unite Sleep Homeostasis and alpha2-Adrenergic Sedation. Curr Biol 29, 3315–3322 e3313.

Mahowald, M.W., Cramer Bornemann, M.A., and Schenck, C.H. (2011). State dissociation, human behavior, and consciousness. Curr Top Med Chem 11, 2392–2402.

McGinley, M.J., Vinck, M., Reimer, J., Batista-Brito, R., Zagha, E., Cadwell, C.R., Tolias, A.S., Cardin, J.A., and McCormick, D.A. (2015). Waking State: Rapid Variations Modulate Neural and Behavioral Responses. Neuron 87, 1143–1161.

McGinty, D., and Szymusiak, R. (2001). Brain structures and mechanisms involved in the generation of NREM sleep: focus on the preoptic hypothalamus. Sleep Med Rev 5, 323–342.

McKinley, M.J., Yao, S.T., Uschakov, A., McAllen, R.M., Rundgren, M., and Martelli, D. (2015). The median preoptic nucleus: front and centre for the regulation of body fluid, sodium, temperature, sleep and cardiovascular homeostasis. Acta Physiol (Oxf) 214, 8–32.

Mochizuki, T., Crocker, A., McCormack, S., Yanagisawa, M., Sakurai, T., and Scammell, T.E. (2004). Behavioral state instability in orexin knock-out mice. J Neurosci 24, 6291–6300.

Modirrousta, M., Mainville, L., and Jones, B.E. (2004). Gabaergic neurons with alpha2-adrenergic receptors in basal forebrain and preoptic area express c-Fos during sleep. Neuroscience 129, 803–810.

Moffitt, J.R., Bambah-Mukku, D., Eichhorn, S.W., Vaughn, E., Shekhar, K., Perez, J.D., Rubinstein, N.D., Hao, J., Regev, A., Dulac, C., et al. (2018). Molecular, spatial, and functional single-cell profiling of the hypothalamic preoptic region. Science 362.

Muheim, C.M., Spinnler, A., Sartorius, T., Durr, R., Huber, R., Kabagema, C., Ruth, P., and Brown, S.A. (2019). Dynamic- and Frequency-Specific Regulation of Sleep Oscillations by Cortical Potassium Channels. Curr Biol 29, 2983–2992 e2983.

Nauta, W.J. (1946). Hypothalamic regulation of sleep in rats; an experimental study. J Neurophysiol 9, 285–316.

Nieh, E.H., Vander Weele, C.M., Matthews, G.A., Presbrey, K.N., Wichmann, R., Leppla, C.A., Izadmehr, E.M., and Tye, K.M. (2016). Inhibitory Input from the Lateral Hypothalamus to the Ventral Tegmental Area Disinhibits Dopamine Neurons and Promotes Behavioral Activation. Neuron 90, 1286–1298.

Noya, S.B., Colameo, D., Bruning, F., Spinnler, A., Mircsof, D., Opitz, L., Mann, M., Tyagarajan, S.K., Robles, M.S., and Brown, S.A. (2019). The forebrain synaptic transcriptome is organized by clocks but its proteome is driven by sleep. Science 366.

Oishi, Y., Xu, Q., Wang, L., Zhang, B.J., Takahashi, K., Takata, Y., Luo, Y.J., Cherasse, Y., Schiffmann, S.N., de Kerchove d’Exaerde, A., et al. (2017). Slow-wave sleep is controlled by a subset of nucleus accumbens core neurons in mice. Nat Commun 8, 734.

Owen, S.F., Liu, M.H., and Kreitzer, A.C. (2019). Thermal constraints on in vivo optogenetic manipulations. Nat Neurosci 22, 1061–1065.

Paxinos, G., and Franklin, K.B.J. (2001). The mouse brain in stereotaxic coordinates, 2nd edn (San Diego: Academic Press).

Sallanon, M., Denoyer, M., Kitahama, K., Aubert, C., Gay, N., and Jouvet, M. (1989). Long-lasting insomnia induced by preoptic neuron lesions and its transient reversal by muscimol injection into the posterior hypothalamus in the cat. Neuroscience 32, 669–683.

Saper, C.B., Chou, T.C., and Scammell, T.E. (2001). The sleep switch: hypothalamic control of sleep and wakefulness. Trends in Neurosciences 24, 726–731.

Saper, C.B., Fuller, P.M., Pedersen, N.P., Lu, J., and Scammell, T.E. (2010). Sleep state switching. Neuron 68, 1023–1042.

Sherin, J.E., Shiromani, P.J., McCarley, R.W., and Saper, C.B. (1996). Activation of ventrolateral preoptic neurons during sleep. Science 271, 216–219.

Shi, S., and Ueda, H.R. (2018). Ca(2+) -Dependent Hyperpolarization Pathways in Sleep Homeostasis and Mental Disorders. Bioessays 40.

Suzuki, A., Sinton, C.M., Greene, R.W., and Yanagisawa, M. (2013). Behavioral and biochemical dissociation of arousal and homeostatic sleep need influenced by prior wakeful experience in mice. Proc Natl Acad Sci U S A 110, 10288–10293.

Szymusiak, R., and McGinty, D. (2008). Hypothalamic regulation of sleep and arousal. Ann N Y Acad Sci 1129, 275–286.

Tatsuki, F., Sunagawa, G.A., Shi, S., Susaki, E.A., Yukinaga, H., Perrin, D., Sumiyama, K., Ukai-Tadenuma, M., Fujishima, H., Ohno, R., et al. (2016). Involvement of Ca(2+)-Dependent Hyperpolarization in Sleep Duration in Mammals. Neuron 90, 70–85.

Tyssowski, K.M., and Gray, J.M. (2019). Blue Light Increases Neuronal Activity-Regulated Gene Expression in the Absence of Optogenetic Proteins. eNeuro 6.

Ungurean, G., van der Meij, J., Rattenborg, N.C., and Lesku, J.A. (2020). Evolution and plasticity of sleep. Current Opinion in Physiology 15, 111–119.

Vassalli, A., and Franken, P. (2017). Hypocretin (orexin) is critical in sustaining theta/gamma-rich waking behaviors that drive sleep need. Proc Natl Acad Sci U S A 114, E5464–E5473.

Vyazovskiy, V.V., and Harris, K.D. (2013). Sleep and the single neuron: the role of global slow oscillations in individual cell rest. Nat Rev Neurosci 14, 443–451.

Vyazovskiy, V.V., and Tobler, I. (2005). Theta activity in the waking EEG is a marker of sleep propensity in the rat. Brain Res 1050, 64–71.

Weber, F., Hoang Do, J.P., Chung, S., Beier, K.T., Bikov, M., Saffari Doost, M., and Dan, Y. (2018). Regulation of REM and Non-REM Sleep by Periaqueductal GABAergic Neurons. Nat Commun 9, 354.

Williams, J.A., and Naidoo, N. (2020). Sleep and Cellular Stress. Curr Opin Physiol 15, 104–110.

Wu, Z., Autry, A.E., Bergan, J.F., Watabe-Uchida, M., and Dulac, C.G. (2014). Galanin neurons in the medial preoptic area govern parental behaviour. Nature 509, 325–330.

Zhang, Z., Ferretti, V., Guntan, I., Moro, A., Steinberg, E.A., Ye, Z., Zecharia, A.Y., Yu, X., Vyssotski, A.L., Brickley, S.G., et al. (2015). Neuronal ensembles sufficient for recovery sleep and the sedative actions of alpha2 adrenergic agonists. Nat Neurosci 18, 553–561.

Zhao, Z.D., Yang, W.Z., Gao, C., Fu, X., Zhang, W., Zhou, Q., Chen, W., Ni, X., Lin, J.K., Yang, J., et al. (2017). A hypothalamic circuit that controls body temperature. Proc Natl Acad Sci U S A 114, 2042–2047.

Zhong, P., Zhang, Z., Barger, Z., Ma, C., Liu, D., Ding, X., and Dan, Y. (2019). Control of Non-REM Sleep by Midbrain Neurotensinergic Neurons. Neuron 104, 795–809 e796.

